# Diverse viruses infect nitrifying archaea and bacteria communities in soil

**DOI:** 10.1101/2023.12.02.569724

**Authors:** Sungeun Lee, Christina Hazard, Graeme W. Nicol

## Abstract

Soil virus communities are diverse and dynamic but contributions to specific processes, such as nitrification, are largely uncharacterised. Chemolithoautotrophic nitrifiers perform this essential component of the nitrogen cycle and are established model groups for linking phylogeny, evolution and ecophysiology due to limited taxonomic and functional diversity. Ammonia-oxidising bacteria (AOB) dominate the first step of ammonia oxidation at high supply rates, with ammonia-oxidising archaea (AOA) and complete ammonia-oxidising *Nitrospira* (comammox) often active at lower supply rates or when AOB are inactive, and nitrite-oxidising bacteria (NOB) completing canonical nitrification. Here, the diversity and genome content of dsDNA viruses infecting different nitrifier groups were characterised after *in situ* enrichment via differential host inhibition, a selective approach that alleviates competition for non-inhibited populations to determine relative activity. Microcosms were incubated with urea to stimulate nitrification and amended with 1-octyne or 3,4-dimethylpyrazole phosphate (AOB inhibited), acetylene (all ammonia oxidisers inhibited), or no inhibitor (AOB stimulated), and virus-targeted metagenomes characterised using databases of host genomes, reference (pro)viruses and hallmark genes. Increases in the relative abundance of nitrifier host groups were consistent with predicted inhibition profiles and concomitant with increases in the relative abundance of their viruses, represented by 200 viral operational taxonomic units. These included 61 high-quality/complete virus genomes 35-173 kb in length and possessing minimal similarity to validated families. Most AOA viruses were placed within a unique lineage and viromes were enriched in AOA multicopper oxidase genes. These findings demonstrate that focussed incubation studies facilitate characterisation of host-virus interactions associated with specific functional processes.

## Introduction

Nitrification is the microbially-mediated sequential oxidation of ammonia (NH_3_) to nitrite (NO_2_^-^) and nitrate (NO_3_^-^) [1]. While it is an essential component of the global nitrogen (N) cycle, linking the most reduced and oxidised forms of N, it also contributes to reduced N use efficiency (NUI) in agroecosystem soils [2], whereby the majority of applied fertilisers are lost before assimilation by crop plant roots [3]. This leads to the loss of mobile NO_3_^-^ via leaching, directly produces the greenhouse gas N_2_O via NH_3_ oxidation, and indirectly contributes to elevated N_2_O production via provision of nitrogen oxide substrates for facultative denitrifying microorganisms under conditions of low oxygen or anoxia [4].

Nitrification in soil is typically dominated by specialised groups of chemolithoautotrophic ammonia-oxidising microorganisms (AOM) and nitrite-oxidising bacteria (NOB) that both derive energy from oxidising inorganic N to fix inorganic carbon (C). Aerobic soil AOM comprise three groups: canonical ammonia-oxidising archaea (AOA) of the families *Nitrososphaeraceae* and *Nitrosopumilaceae* of the class *Nitrososphaera* [5]; canonical ammonia-oxidising bacteria (AOB) of the genera *Nitrosomonas*, *Nitrosospira* and *Nitrosococcus* (with *Nitrosomonas* and *Nitrosospira* typically dominating in agricultural soils [6]); and complete ammonia-oxidizing bacteria of the genus *Nitrospira* (comammox) which oxidise NH_3_ through to NO_3_^-^ in a single cell [7,8]. Canonical NOB in soil include representatives of the genera *Nitrospira*, *Nitrobacter, Nitrococcus* and *Nitrotoga* [9], with *Nitrobacter* and *Nitrospira* typically dominating in agricultural soils [6].

While AOB were thought to be the only group of AOM found in soil for over a century [10], since the discovery of AOA and comammox, the use of compounds which inhibit specific AO groups have been widely used to examine niche differentiation *in situ* [11]. Application of inhibitors alleviates competition for NH_3_ by non-inhibited groups and their relative contribution or ecophysiological features inferred [12–14]. These include compounds tested specifically as inhibitors for all autotrophic ammonia oxidisers to measure heterotrophic nitrification activity (e.g. acetylene [15]), AOB-specific inhibitors (e.g. 1-octyne[16]) or AOA-specific inhibitors (e.g. 2-pheny l-4,4,5,5-tetramethylimidazoline-1-oxyl 3-oxide (PTIO) [17]). Other compounds have been used in agriculture to increase NUI after fertiliser application, but laboratory culture or soil molecular analyses can demonstrate preferential inhibition of one AOM group. For example, AOB have higher sensitivity to 3,4-dimethylpyrazole phosphate (DMPP) [18] or allylthiourea [19,20] compared to AOA. Differential inhibition of soil AOB has demonstrated they dominate activity and outcompete AOA at high supply rates of NH_4_^+^ [4], producing double the yield of N_2_O compared to AOA [11], but AOA can utilise high supply rates of NH_4_^+^ when AOB are inhibited specifically [12].

The impact of virus activity on specific functional processes in soil, including N cycling, are generally uncharacterised. While augmenting viral loads have been demonstrated to alter inorganic N content in constructed systems [21], decreases in the abundance of growing soil nitrifier populations, or variation in N fluxes that could be attributed to viral predation via ‘kill-the-winner’ dynamics [22] are typically not observed in soils. However, total prokaryote community host-virus interactions are highly dynamic in soil and AOA viruses have been demonstrated to be active during nitrification using ^13^C-tracing experiments [23]. Comparatively little is known about the diversity of viruses infecting AOM and NOB compared to other cultivated taxonomic groups that are ubiquitously distributed in soil. This may be in part due to cultivated strains of AOM and NOB only forming (micro)colonies on solidified media without confluent growth [24–26], limiting the use standard plaque assays for isolating viruses that are reproducing via the lytic cycle. Nevertheless, spindle-shaped viruses, which represent an archaea host-specific morphology [27], were isolated from marine water after infection of AOA *Nitrosospumilus* strains [28], and the first cultivated lytic virus of AOB (ɸNF-1) was also recently isolated from wastewater [29], infecting different strains of *Nitrosomonas*. Cultivated AOA, AOB and NOB genomes contain integrated proviruses (e.g. [30–34], CRISPR-Cas systems (e.g. [35–37]) and a variety of other viral defence mechanisms [38], indicating dynamic interactions with viruses. However, due to a lack of previously characterised viruses, the taxonomy of those infecting soil nitrifiers is unknown.

The overall aim of this study was to characterise soil virus populations infecting different nitrifier groups in urea-stimulated microcosms also amended with a series of inhibitors. It was hypothesized that nitrification would result in increases in the relative abundance of viruses infecting nitrifiers demonstrating their activity in soil, and targeted alleviation of competition for NH_3_ would enrich viruses infecting uninhibited populations, providing a mechanism for characterising AOB, AOA, NOB and comammox viruses under different growth conditions.

## Material and methods

### Soil microcosms

Soil was sampled in February 2022 from the upper 10 cm of a sandy loam agricultural soil (Rozier Abbey urban farm, Ecully, France; latitude/longitude 45.777/4.788). The soil is under crop rotation and was previously used for cultivating green beans (*Phaseolus vulgaris*). Soil was sampled from three 1 m^2^ quadrats separated by 5 m intervals along a transect using a surface-sterilised (70% ethanol (v/v)) trowel to generate triplicate samples which were homogenised individually by sieving (2 mm mesh) and stored at 4°C prior to establishing microcosms and physicochemical analyses. Water content was determined by mass loss after drying at 105°C for 24 h. Soil samples had a pH in water of 7.2 (±0.1 s.e.) (2:1 water:soil ratio), 5.1% (±0.1%) total organic matter content (loss on ignition; 450°C for 24 h) and 2.4% (±0.2%) and 0.17% (±0.01%) total C and N, respectively (Carlo Erba NC 2500 elemental analyser). Soil microcosms were established in 120 ml serum bottles with 30 g soil (dry weight (_dw_) equivalent) with an initial 18% (w/w) water content. Soil microcosms were pre-incubated at 25°C for 5 days before the addition of 200 µg urea-N g^-1^ soil_dw_ (or water only (control)) together with individual inhibitors; 1-octyne (0.03% (v/v) headspace concentration) to inhibit AOB [16], 3,4-dimethylpyrazole phosphate (DMPP) (0.5% of applied N) to inhibit AOB [39], acetylene (0.1% (v/v) headspace concentration) to inhibit all ammonia oxidisers [40] or no inhibitor (urea only), with soil in all microcosms having a 20% (w/w) water content after amendments. In this soil, all urea applied at this concentration is hydrolysed to NH_4_^+^ within 24 h (data not shown). All microcosms were opened and aerated every five days to maintain aerobic conditions before re-establishing gaseous inhibitor concentrations. Microcosms were destructively sampled in triplicate after 0, 5, 10, 15, 20, 25 and 30 days of incubation with a further 200 or 100 μg urea-N g^−1^ soil_dw_ added to prevent NH_3_ limitation when concentrations decreased below 50 or 100 μg NH_4_^+^-N g^−1^ soil_dw_, respectively. Microcosms not receiving additional urea were amended with the same volume of water (0.3 ml) resulting in increases in water content to 21 and 22% (w/w) at day 10 and 20, respectively, for all microcosms. Virus DNA extractions (day 0 and 30) and determination of inorganic N concentrations (all time-points) were performed immediately after sampling with a portion of soil (∼5 g) archived at -20°C for total soil DNA extraction. NH_4_^+^, NO_2_^-^ and NO_3_^-^ concentrations were determined using standard colorimetric assays [12]. Recovery of soil virus particles was based on the protocol of Trubl et al. [41] and virus DNA subsequently extracted using a CTAB/SDS/proteinase K protocol modified from Casas and Rower (2017), both with modifications as described by Lee et al. [42]. Total genomic DNA was extracted from 0.5 g soil using the CTAB/phosphate buffer/phenol-chloroform-isoamyl alcohol bead-beating protocol [43].

### *16S* rRNA gene amplicon sequencing and bioinformatic analysis

PCR amplification of prokaryote 16S rRNA genes was performed using primers 515F and 806R [44] with Illumina adapters. PCR was performed in 25 µl reactions using Invitrogen Platinum Taq DNA polymerase (Thermo Fisher), 0.5 ul of forward and reverse primers (0.2 µM final concentration) and 2 µl of template DNA (2 ng total). Thermocycling conditions were 95°C for 3 min; 30 cycles of 95°C for 30 s, 55°C for 30 s, 72°C for 30 s; and 72°C for 5 min. Amplicons were bead purified using Agencourt AMPure XP (Beckman Coulter) before indexing PCR. Indexed amplicons were bead purified followed by photometric quantification using a μDrop plate (Thermo Fisher). Equimolar concentrations were pooled and quality-controlled using a Bioanalyzer with DNA 1000 Kit (Agilent) before sequencing using the MiSeq platform (Illumina) with Reagent Kit v2 (500-cycles). Sequence data were analysed using the DADA2 tool (v.1.1.6) based on an amplicon sequence variant (ASV)-based pipeline [45]. Taxonomic affiliation and count tables of ASVs were generated using assignTaxonomy function against the SILVA database (release 138.1) [46]. *amoA* gene-defined designations of AOA lineages [5] were defined for 16S rRNA gene ASVs using the pipeline of Wang et al. [47].

### Virome sequencing and vOTU prediction

Virome DNA was sequenced using the NovaSeq platform (Illumina) at IntegraGen (Paris, France). Raw reads were processed using MetaWrap read_qc module [48] as described in Lee et al. [49]. Co-assembly of quality-controlled reads was performed using Megahit v1.1.2 [50], resulting in 56,617 10-kb contigs. Contigs of viral origin were predicted from those ≥10 kb using VirSorter [51], VirSorter 2.0 [52] and DeepVirFinder [53], geNomad [54] used to confirm a viral origin for conspicuously large contigs, and quality and completeness assessed using CheckV (checkv-db-v1.5) [55] and VIBRANT v1.2.0 [56]. Viral contigs were clustered into viral operational taxonomic units (vOTU) using BLASTn v.2.11.0 with a global identity ≥95% and coverage ≥85% in accordance with recommended standards [57]. Gene prediction was performed using Prodigal v2.6.3 with meta option [58] and initial annotation performed using Diamond BLASTp v0.8.36 with the NCBI nr database release 244 [59]. Gene prediction and annotation were additionally performed with the VIPTree server using the tools GeneMarkS [60] and GHOSTX [61] with the NCBI/nr database and hmmsearch function in HMMER 3.3.2 [62] with VOG HMM database (http://vogdb.org). Manual curation (identifying structural and replication viral hallmark genes, depletion in annotation genes, enrichment of hypotheticals) was also performed for nitrifier host-predicted vOTUs. Lysogenic potential was predicted using VIBRANT [56] plus manual curation from annotation tables. The abundance of vOTUs in Rozier and soil samples from other studies was estimated using BBMap v38.96 [63] and BamM v1.7.3 [64]. vOTU detection in a soil sample was inferred from a detection threshold of ≥85% of contig length with ≥1x read recruitment at ≥95% average nucleotide identity [57]. For detection of viruses in other soils sharing genomic content (but not interpreted as detection of the same virus), a lower threshold was used of ≥10% contig length at ≥90% average nucleotide identity. Normalised relative abundance was expressed as reads per kilobase per million (RPKM) mapped reads. Heatmap representation of relative abundance was produced using the pheatmap R package [65] in R v4.2.2. Linear regression between virus and host abundance was performed using ggscatter and stat_cor function with the ggpubr R package [66].

### Host prediction and analysis

The use of CRISPR array spacer matching for predicting hosts of viruses was evaluated using 487 CRISPR arrays predicted from predicted 550 nitrifier (AOA, AOB and NOB/comammox) genomes selected from the GTDB database. No matches were identified with less than 2 mismatches and linkages were subsequently predicted using a shared gene-based approach [49,67]. vOTUs from viruses potentially infecting autotrophic nitrifiers were first screened with “best hit” (amino acid identity >30%, e-value <10^−5^, bit score >50, and query coverage >70%) matches to a minimum of three homologs in nitrifier host genomes and further assessed using the automated host prediction tool iPHoP v1.1.0 tool [68]. vOTUs were then compared against a reference database of nitrifier virus genes derived from i) 2,399 provirus sequences identified in nitrifier host genomes selected from the GTDB database [69] and ii) 1,078 predicted complete or high-quality (predicted ≥90% complete) contigs from the IMG-VR v4 database [70] predicted to infect nitrifier taxa. Provirus regions were identified from GTDB nitrifier genomes using PhageBoost v0.1.7 [71], VIBRANT v1.2.0 [56] and VirSorter [51]. Gene prediction and annotation were then performed as described previously. A nitrifier virus hallmark gene database of structural and replication genes was generated from annotated genes in CheckV quality-passed provirus sequences [55].

Gene sharing network analysis between Rozier nitrifier virus vOTUs and reference sequences was performed using vConTACT2.0 [72]. Whole genome comparisons using tBLASTx scores were performed using ViPTree [73]. Using predicted high-quality/complete vOTUs only, the diversity of potential virus families infecting each nitrifier group was also assessed with ViPTree using a proposed threshold cut-off of 0.05 that correlates with an observed demarcation for previously validated virus families [74]. Genome maps of individual nitrifier virus families with more than one representative were made using EasyFig genome comparison visualiser v2.2.3 [75] with BLAST output files provided from the ViPTree server [73].

### Phylogenetic analysis of individual genes

Phylogenetic analysis of terminase large subunit (TerL) genes from nitrifier vOTU and RefSeq release 218 [76] viruses was performed using an alignment generated with MAFFT v7.505 using the einsi algorithm [77]. Ambiguous aligned regions were removed using the TrimAI v1.4 tool with gappyout option [78] and phylogeny calculated using IQ-TREE v.2.2.5 [79] with automatic substitution model selection. Phylogenetic analysis of auxiliary metabolic genes (AMGs) was performed using alignments generated using MUSCLE [80] and constructed using unambiguously aligned positions with PhyML and automatic model selection.

## Results

### Nitrification in microcosms amended with urea and nitrification inhibitors

Net nitrification was highest in uninhibited urea-amended microcosms with 554 (±62.4 s.e.) µg NO_3_^-^-N g^-1^ soil_dw_ produced after 30 days. The addition of DMPP, 1-octyne or acetylene resulted in inhibition of NH_3_ oxidation with expected decreases in nitrification rates. Acetylene completely inhibited nitrification confirming heterotrophic nitrification did not occur [6]. Nitrate production was lower with AOB inhibitors, but DMPP had a significantly greater effect than 1-octyne (*p*=0.002) with net production of 221 (±17.5) μg NO_3_^-^-N g^-1^ soil_dw_ and 426 (±58.5). While this could indicate non-AOB AOM were also inhibited, it is possible that some AOB populations were insensitive to 1-octyne. Complete inhibition by acetylene in these well-aerated microcosms with relatively low water content demonstrated that gas diffusion was not an issue, and further incubations with higher concentrations of 1-octyne did not increase inhibition rates (data not shown).

### Selective enrichment of specific nitrifier communities after differential inhibition

Changes in the relative abundance of 16S rRNA genes from populations belonging to the genera *Nitrosomonas* and *Nitrosospira* (corresponding to AOB), the class *Nitrososphaeria* (corresponding to AOA), and *Nitrospira*and *Nitrobacter* (corresponding to NOB) were determined in amplicon sequence libraries from day 0 and 30 samples. Sequences associated with other bacterial nitrifier groups (e.g. *Nitrosococcus* AOB or *Nitrotoga* NOB) were either absent or represented <0.1% of AOB or NOB 16S rRNA sequences, respectively.

Relative to all prokaryote 16S rRNA gene amplicons, AOB 16S rRNA gene abundance increased significantly (*p* < 0.05) from 0.07(±0.04)% at day 0 to 0.6(±0.3)% (8-fold increase) and 1.5(±0.2)% (22-fold increase) at day 30 in 1-octyne and uninhibited urea-microcosms, respectively (Fig. 1B), with all ASVs affiliated with the genus *Nitrosospira*. AOB abundance did not increase in microcosms amended with acetylene or DMPP, indicating that DMPP was a more effective AOB-specific inhibitor than 1-octyne in this soil. AOA relative abundance increased significantly in all microcosms (except for acetylene). AOA relative abundance increase was greatest in microcosms where AOB were partially or fully inhibited, with 2.8, 2.4 and 1.6-fold increases from 1.7(±0.1)% to 4.8(±0.4)%, 4.1(±0.5)% and 2.7(±0.2)% of all 16S rRNA genes in microcosms amended with DMPP, 1-octyne or urea-only, respectively. Increases in relative abundance were associated with ASVs associated with the *amoA*-defined lineages NS-ζ-2 (which includes *Candidatus* Nitrosocosmicus cultivated representatives) and NS-δ-1 (with no cultured representatives) (Fig. S1). *Nitrospira* 16S rRNA gene relative abundance increased significantly in microcosms amended with 1-octyne only from 1.7(±0.2) to 3.6(±0.2)% (2.1-fold increase) with all ASVs affiliated with the genus *Nitrospira*. Phylogenetic analysis of 16S rRNA genes does not provide a framework for differentiating canonical NOB vs comammox *Nitrospira* but an increase in relative abundance when AOB were inhibited suggests a proportion were oxidising NH_3_. *Nitrobacter* 16S rRNA gene abundance was less than 10% of *Nitrospira* 16S rRNA genes in all samples and no significant changes in relative abundance were observed under any amendment (data not shown).

**Fig. 1.**
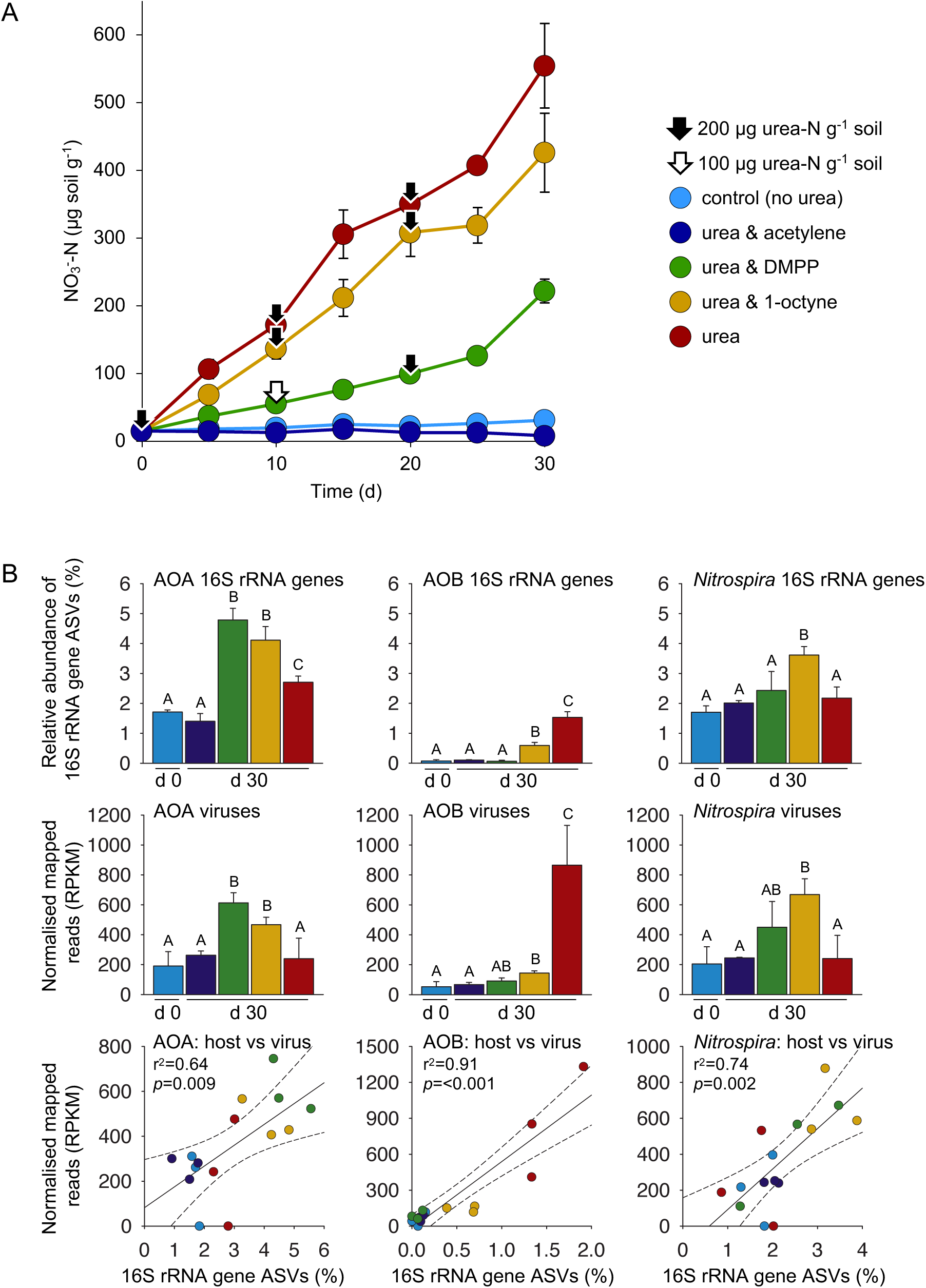
Nitrification activity and abundance of nitrifier hosts and viruses in differentially-inhibited soil microcosms amended with urea. **A** NO_3_^-^ concentrations in soil microcosms amended with 200 µg urea-N g^-1^ soil and NH_3_ oxidation inhibitors acetylene, DMPP, 1-octyne or control (no inhibitor). An additional 100 or 200 µg urea-N g^-1^ was added when NH_4_^+^ concentrations were below 100 or 50 µg NH_4_^+^-N g^-1^, respectively, to prevent NH_3_ limitation. **B** Relative abundance (%) of 16S rRNA ASVs of AOA, AOB and *Nitrospira* in total prokaryote 16S rRNA amplicon libraries after 0 and 30 days incubation (d 0 and d 30, respectively). Samples with different letters indicate significant differences (*p* < 0.05, two-sample Student’s t-test or Welsch’s t-test when variances were not homogenous). **C** Relative abundance (RPKM) of reads mapped to contigs derived from genomes of viruses predicted to infect AOA, AOB and *Nitrospira* hosts. Samples with different letters indicate significant differences (p<0.05). **D** Correlation between paired host and virus relative abundance in individual samples for AOB, AOA and *Nitrospira*. Dotted lines denote 95% confidence intervals. For all panels, error bars represent the standard error of the mean of triplicate samples each derived from an individual field replicate.

### Selective enrichment of nitrifier viruses after differential inhibition

Viromes were prepared for day 0 and 30 samples with an average of 125 million (range 77-218 million) quality-filtered reads (Table S1). Reads from 16S rRNA gene sequences indicating cellular DNA contamination represented only 0.002% of reads and was comparable with other virome-based studies (e.g. [81]). After co-assembly, contigs ≥10 kb representing 17,817 viral operational taxonomic units (vOTU) were identified. A total of 16.3% were linked to predicted hosts including *Proteobacteria* (11.6%), *Actinobacteria* (1.8%) and *Bacteroidetes* (1.0%) and consistent with the high prevalence of *Proteobacteria* (16.7%), *Actinobacteria* (8.5%) and *Bacteroidetes* (7.9%) in 16S rRNA gene amplicon sequencing analysis. Two hundred vOTUs (1.1%) were predicted to represent viruses infecting nitrifiers (AOA (n=39), AOB (n=62), comammox and canonical NOB (n=99)). These were also analysed using the host-prediction framework iPHoP and while 67 (33.5%) vOTUs were predicted to have a nitrifier host using at least one of the six individual classifiers implemented in iPHoP, only 15 (7.5%) had a host predicted with high confidence (iPHoP score >90) of which only 5 were nitrifiers. However, the majority of vOTUs (67.5%) were placed in gene-sharing networks with reference nitrifier viruses in vConTACT analyses and 74 (37%) contained genes possessing identity (cut-off values) with reference nitrifier virus hallmark genes (described in further detail below).

Predicted nitrifier vOTU relative abundance was determined by read-mapping. Increases in viral abundance after 30 days incubation were fully concomitant with changes in host abundance and significant increases observed for AOB-infecting viruses in 1-octyne and urea (no inhibitor) microcosms only, for AOA-infecting viruses in DMPP and 1-octyne microcosms only, and for *Nitrospira*-infecting viruses in 1-octyne microcosms only. The r^2^ of all correlations between host and virus relative abundance ranged from 0.64 to 0.91 and all were significant (*p* ≤ 0.001-0.009) (Fig. 1B).

### Characterisation of AOA virus diversity

Reference AOA virus genomes included sequences from integrated proviruses (n=1,101) identified in 212 *Nitrososphaeria* genomes accessed from GTDB and 707 high-quality or complete metagenome-derived virus contigs from the IMG-VR. Of these, 121 (6.6%) reference sequences were in gene-sharing networks with 38 (97.4%) AOA virus sequences from Rozier soils (Figure 2A). Twenty-one vOTUs recovered from a Scottish agricultural soil (‘Craibstone’) and predicted to infect AOA [23] were also placed within the network. Integrated proviruses infecting the same taxonomic family grouped together, including those infecting AOA families *Nitrosopumilaceae*, *Nitrososphaeraceae* and *Nitrosocaldaceae* plus other non-AOA *Nitrososphaeria*. Rozier and Craibstone virus contigs clustered together with the majority also linked to reference viruses infecting the *Nitrososphaeraceae* (including representatives of the genera *Nitrososphaera*, *Nitrosocosmicus*, TH5896) but also *Nitrosopumilaceae* and *Nitrosocaldaceae* hosts. Three contigs were also linked to proviruses of non-AOA *Nitrosophaeria* found in soil, such as the lineage UBA183/Group 1.1c/ *Gagatemarchaeaceae* [69,82,83].

**Fig. 2.**
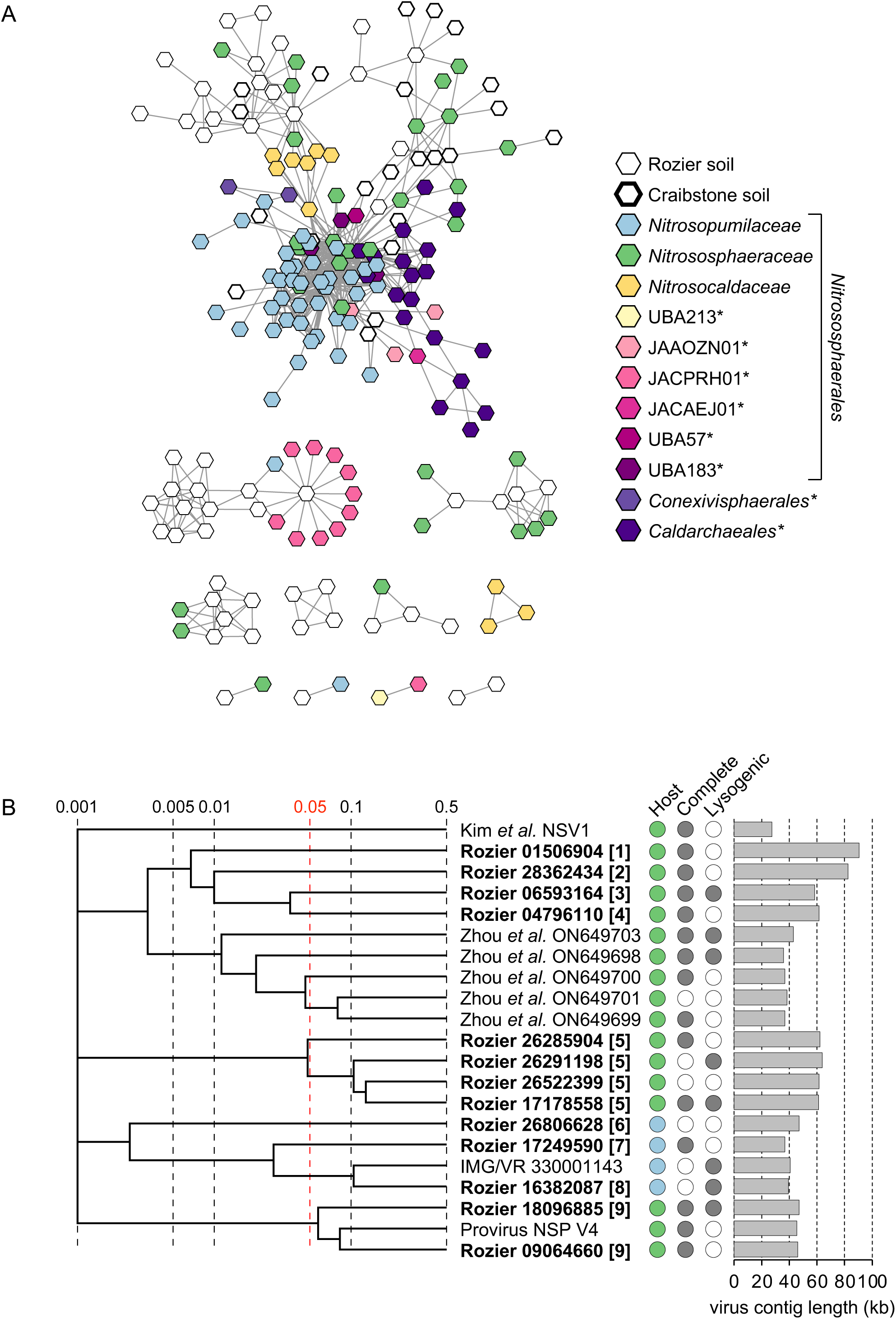
Diversity and relatedness of contigs from predicted AOA-infecting virus genomes. **A** Gene-sharing network analysis of all virus contigs ≥10 kb from this study (Rozier soil) and our previous study (Craibstone soil; [23]) associated with hosts of the class *Nitrososphaeria*. Reference virus sequences are provirus sequences extracted from *Nitrososphaeria* host genomes and complete or high-quality genomes from the IMG/VR database with a predicted *Nitrososphaeria* host. Non-AOA (selected families within the *Nitrososphaerales* or *Caldarchaeales* and *Conexivisphaerales* orders) are denoted with *. **B** Proteomic tree showing genome-wide sequence similarities between complete or high-quality AOA viral contigs from Rozier soil (in bold with eight-figure NCBI contig reference) and reference sequences. Colour coding for predicted host follows the key in panel A, and genome completeness and prediction of lysogeny is denoted with filled circles. Values at dotted lines represent a distance metric based on normalized tBLASTx scores with 0.05 (in red) an estimated threshold for grouping viruses within the same family. Each number in a square bracket denotes an individual putative virus family.

Thirteen virus contigs were predicted to be complete (10) or high-quality (3), ranged in size from 36.9 to 90.5 kb in size (Figure 2B; Figure S2) and five were predicted to be capable of lysogeny. Consistent with correlation in virus and host codon usage [84] and the low GC mol% of AOA genomes [23], AOA virus vOTUs had a mean GC mol% of 43.7% and lower than the mean of 54.7% for all Rozier virus contigs. Using a criteria of ViPTree scores ≥0.05 that represents an approximate demarcation for individual virus families as proposed by Zhou et al. [74], these represented 10 different putative virus families. Rozier soil AOA viruses only possessed identity with other high-quality virus sequences from integrated AOA proviruses or metagenome-derived sequences. No identity was observed with either RefSeq viruses or marine *Nitrosopumilus* AOA-infecting spindle-shaped marine viruses [28]. Some shared a low level of identity with metagenome-derived virus contigs derived from marine harbour water which were related to proviruses found in soil-derived *Nitrososphaera*, rather than marine-derived *Nitrosopumilus* hosts [74], indicating a potential allochthonous origin in that study.

To further investigate the breadth of AOA-infecting virus diversity, phylogenetic analysis of large sub-unit terminase (TerL) inferred protein sequences was performed with 1,573 RefSeq-derived sequences (Fig. S3A). While six diverse lineages were observed, the majority were placed in one cluster with AOA provirus-derived TerL sequences. Whole genome analysis also indicated that the majority of contigs are placed within one broad soil-specific lineage including AOA-infecting viruses identified in agricultural soils from Scotland [23] and the USA [85] (Fig. S3B).

To widen the search for viruses potentially infecting AOA, vOTUs containing archaeal host homologs (i.e. not restricted to AOA/*Nitrososphaerales*) were also examined with eight (five complete, three high-quality) belonging to seven different putative families (Fig. S4A), and seven of which were placed in one gene-sharing network with *Nitrososphaeria* reference viruses together with smaller (≥10 kb) incomplete vOTUs (Fig. S4B). This included the identification of contigs with conspicuously larger genomes (191-218 kb) and therefore placed within the “jumbo phage” category [86] (Fig. S4C). While the three largest and related viral genomes (Fig. S4C) possessed up to eight BLASTp ‘best hit’ matches to homologs in nitrifier hosts, the majority of archaea-like genes were to homologs of *Pacearchaeota* genes of the DPANN lineage, representing up to 4% of all genes in one contig. The GC mol% of these large virus genomes were low and comparable to those of AOA-infecting virus and host genomes. However, no prediction of host could be determined with confidence using our manual curation-centric approach nor iPHoP (Table S2).

### Characterisation of AOB virus diversity

AOB vOTUs were compared with reference AOB virus genomes derived from integrated proviruses (n=745) identified in 79 AOB genomes (*Nitrosomonas* and *Nitrosospira* of the *Nitrosomonadaceae*; *Nitrosococcus* of *Nitrosococcaceae*) accessed from GTDB and 298 high-quality or complete metagenome-derived virus contigs with predicted hosts from the IMG-VR. Of these, 124 (11.8%) reference sequences were in gene-sharing networks together with 52 (82.2%) AOB vOTUs from Rozier and Craibstone soils (Figure 3A). The majority of reference virus genomes from *Nitrosomonas* and *Nitrosospira* were linked in one cluster with separation of viruses associated with each genus. The majority of Rozier and Craibstone virus contigs clustered with *Nitrosospira* reference virus genomes in one of two separate clusters in addition to groupings lacking any references sequences.

**Fig. 3.**
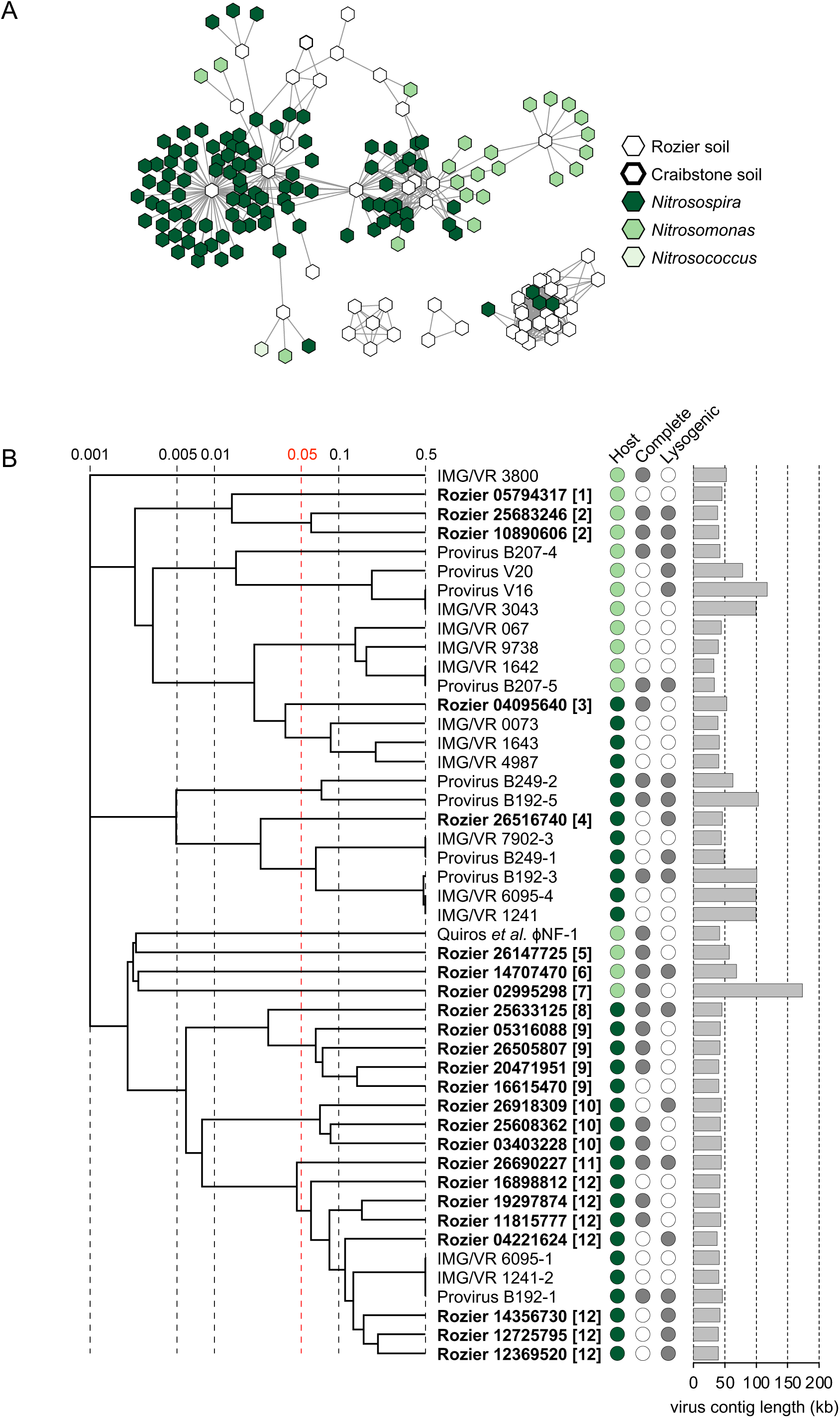
Diversity and relatedness of contigs from predicted AOB-infecting virus genomes. **A** Gene-sharing network analysis of all virus contigs ≥10 kb from this study (Rozier soil) and a previous study (Craibstone soil; [23]) associated with hosts of the genera *Nitrosomonas* and *Nitrosospira*. Reference virus sequences are provirus sequences from host genomes and complete or high-quality genomes from the IMG/VR database with a predicted AOB host. **B** Proteomic tree showing genome-wide sequence similarities between complete or high-quality AOB viral contigs from Rozier soil (in bold with eight-figure NCBI contig reference) and reference sequences. Colour coding for predicted host follows the key in panel A, and genome completeness and prediction of lysogeny is denoted with filled circles. Values at dotted lines represent a distance metric based on normalized tBLASTx scores with 0.05 (in red) an estimated threshold for grouping viruses within the same family. Each number in a square bracket denotes an individual putative virus family.

Twenty-four virus contigs were predicted to be complete (15) or high-quality (9), ranged in size from 37.9 to 173.3 kb in size and 11 (46%) were predicted to be capable of lysogeny (Figure 3B; Figure S5). These represented 11 potentially different families. As with the network analysis of all AOB vOTUs, the majority (18) of Rozier high-quality virus genomes were associated with *Nitrosospira* hosts and grouped with *Nitrosospira* reference sequences in genome-wide comparisons. The six *Nitrosomonas* viruses either grouped with *Nitrosomonas* reference sequences or, at a low level, with *Nitrosomonas*-infecting lytic virus ɸNF-1, recently isolated from wastewater [29], likely reflecting the dominance of *Nitrosospira* rather than *Nitrosomonas* AOB in the soil studied here.

Eight diverse lineages were observed in phylogenetic analysis of Rozier-derived TerL inferred protein sequences with only two lineages, representing 23 (37.0%) Rozier sequences, grouping with reference AOB sequences.

### Characterisation of NOB virus diversity

vOTUs of viruses infecting NOB and potentially comammox *Nitrospira* were compared with reference virus genomes derived from integrated proviruses (n=553) identified in 259 genomes (*Nitrobacter, Nitrococcus*, *Nitrolancea*, *Nitrospina*, *Nitrospirales* (which includes the genus *Nitrospira) and Nitrotoga*) accessed from GTDB and 73 high-quality or complete metagenome-derived virus contigs with predicted hosts from the IMG/VR. Of these, only 18 (2.2%) reference sequences were in gene-sharing networks together with 23 (23.2%) NOB vOTUs from Rozier soils (Figure 3A) and only 5 and 2 vOTUs had direct linkages with *Nitrospira* and *Nitrobacter* reference viruses, respectively, with the majority grouping in networks only with other Rozier soil vOTUs.

Twenty-four virus contigs were predicted to be complete (12) or high-quality (12), ranged in size from 34.5 to 117.8 kb in size and eight (33%) were predicted to be capable of lysogeny. (Figure 4B; Figure S6). These represented 19 potentially different families. As observed for analysis of all vOTUs potentially infecting NOB, there were a lower number of reference virus sequences that shared genome identity.

**Fig. 4.**
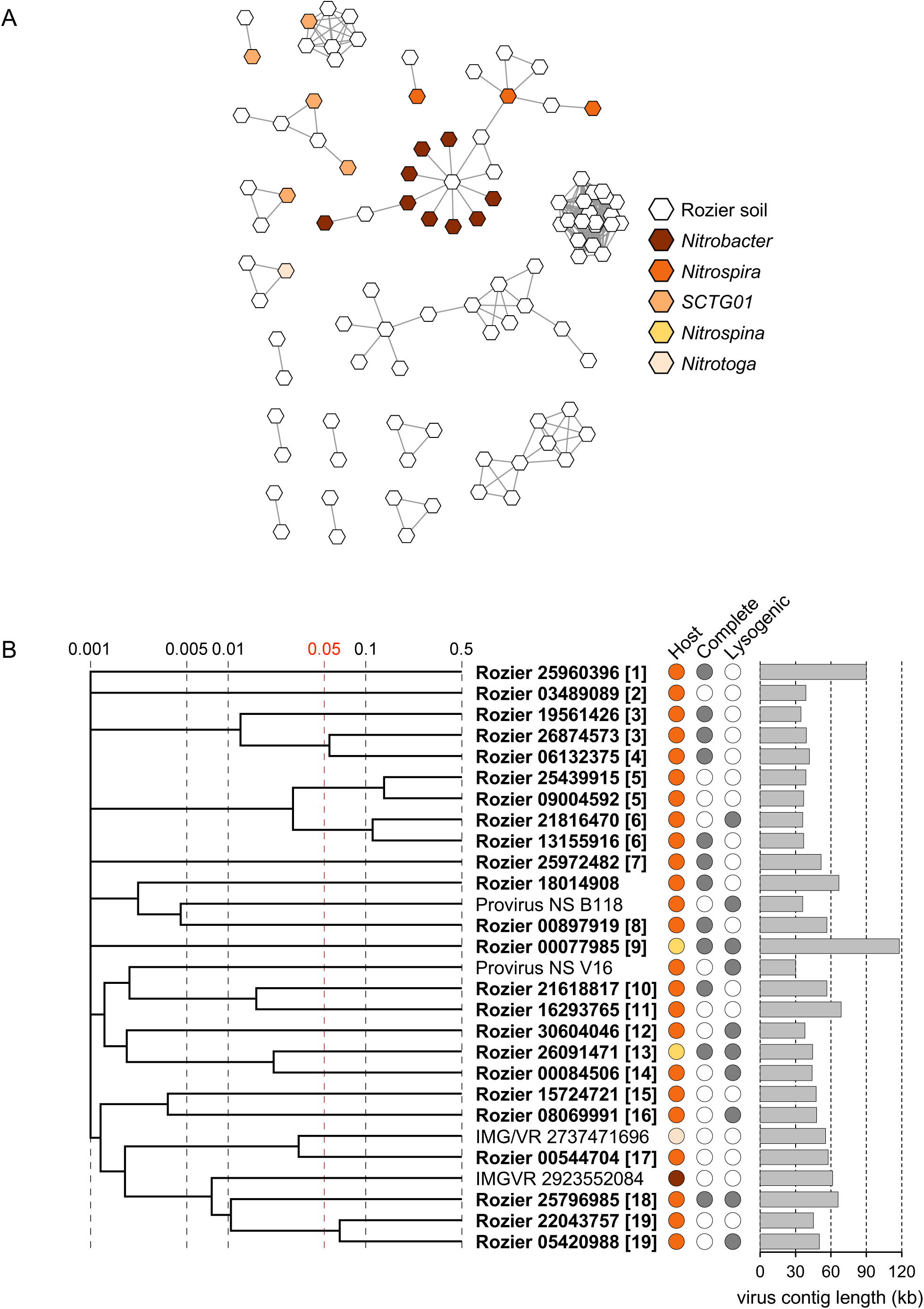
Diversity and relatedness of contigs from predicted NOB-infecting virus genomes. **A** Gene-sharing network analysis of all virus contigs ≥10 kb associated with NOB hosts of different phylogenetic lineages. Reference virus sequences are provirus sequences extracted from host genomes and complete or high-quality genomes from the IMG/VR database with a predicted NOB host. **B** Proteomic tree showing genome-wide sequence similarities between complete or high-quality NOB viral contigs from Rozier soil (in bold with eight-figure NCBI contig reference) and reference sequences. Colour coding for predicted host follows the key in panel A, and genome completeness and prediction of lysogeny is denoted with filled circles. Values at dotted lines represent a distance metric based on normalized tBLASTx scores with 0.05 (in red) an estimated threshold for grouping viruses within the same family. Each number in a square bracket denotes an individual putative virus family.

### Similarity of nitrifier viruses in other soils

The majority of nitrifier vOTUs from this study were not detected in virome datasets from other soils. Using the PIGEON database [87] comprising 266k species-level vOTUs, only 45 (22%) were placed in vConTACT clusters with low-quality (incomplete) vOTUs ≥10 kb in length (Table S3). Forty-two soil viromes representing a range of land use types and soil physicochemical properties (Table S4) were examined using a read-recruitment threshold of 1x coverage ≥85% contig length at 95% identity [57]. However, Rozier-derived vOTUs were not detected in these soils at this threshold and lower threshold of 10% coverage at 95% identity was used for identifying viruses with shared genetic content and resulted in linkage to 59 of 200 Rozier nitrifier vOTUs. While the number ranged from 0 to 20 per soil sample, soil pH had a significant effect (*p =* 1 x 10^-5^) (Fig. S7B) with the highest number linked to viruses in other soils with a similarly neutral pH. These data were also consistent with previous work identifying a relationship between total virus community structures and soil pH [49].

In a recent study of viruses in soils under different long-term nitrogen fertiliser regimens at the West Tennessee Research and Education Center (WTREC), Duan et al. [85] identified vOTUs (≥10 kb) that were predicted to have *Nitrososphaerales*/AOA hosts. While not grouping in our vConTACT analysis, broader level genomic comparisons revealed that they were placed in groupings containing exclusively AOA vOTU representatives from Rozier, Craibstone and WTREC soils (Figure S2b).

### Identifying potential nitrifier-specific auxiliary metabolic genes

Two incomplete contigs (contig 08162653, 14.2 kb; contig 08248335, 39.5 kb) contained homologs of AOA-specific type 1 multicopper oxidases (MCO1) [88] in addition to other genes found in AOA genomes and virus-specific genes (Figure 5AB). The relative abundance of MCO1-encoding AMGs in viromes was further investigated using an assembly-free approach by mapping reads to a database of selected AOA core genes including ammonia monooxygenase sub-units A, B and C (*amoA*, *amoB* and *amoC*), nitrite reductase (*nirK*), MCO1 and type 4 multicopper oxidases (MCO4) (Figure 5C). Consistent with identifying only MCO1 genes in AOA-infecting virus contigs, reads mapped to MCO1 were the most abundant, being 6.7 and 2.6 times more abundant than *amoC* in day 0 and 30 viromes, respectively, which was the second most abundant of the six selected genes.

**Fig. 5.**
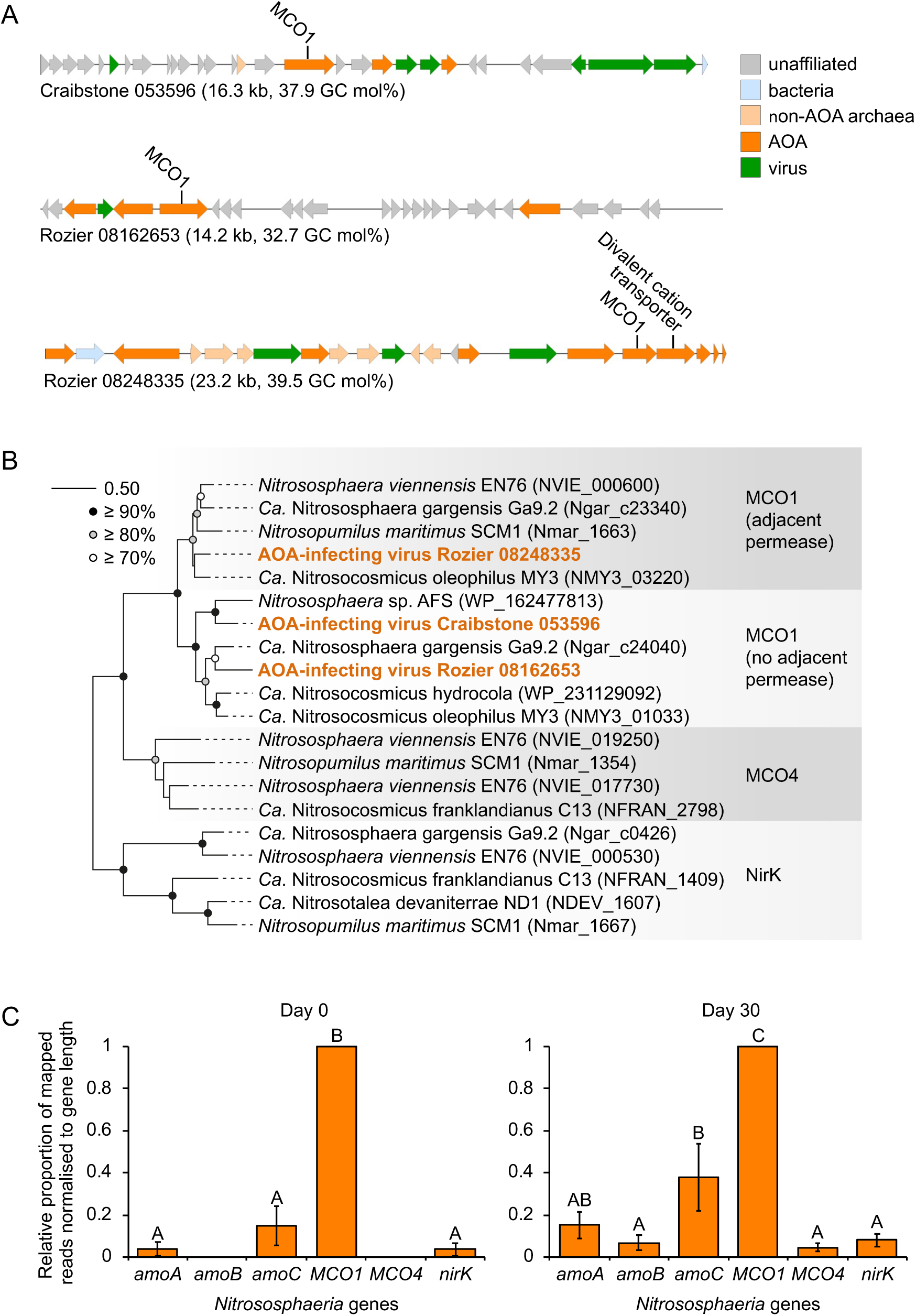
MCO1 genes in AOA-infecting viruses. **A** Genome maps of three predicted virus contigs from two soils (Rozier, this study; Craibstone, Lee et al., [23]) containing MCO1 genes. **B** Maximum likelihood phylogenetic tree describes placement of MCO1 within the AOA MCO/NirK family of cultivated AOA (206 unambiguously aligned positions, LG substitution model with gamma distributed sites). **C** Relative abundance of MCO1 genes in viromes from day 0 and day 30 microcosms compared to other selected AOA-specific core genes. Abundances were normalised by gene length after read mapping. Samples with different letters indicate significant differences (*p* < 0.05, two-sample Student’s t-test or Welsch’s t-test when variances were not homogenous).

Other potential nitrifier-specific AMGs were identified that were not involved in virus structure or replication but specifically shared a highest level of similarity with homologs found in nitrifier genomes. These included a gene encoding an aminotransferase and found in three related medium- or high-quality NOB vOTUs and indicating a role in nitrogen metabolism (Fig. S8). Two AOA virus contigs contained different classes of oxidoreductases also found in multiple AOA genomes and therefore potentially involved in augmenting energy conservation during infection. Three jumbo virus genomes (>200 kb) of putative non-AOA archaea-infecting vOTUs all possessed *phoH* (Fig. S4), an AMG implicated in altering host phosphate metabolism and prevalent in marine viruses but also widely distributed in soil viruses [89].

## Discussion

Nitrifier populations may represent an ideal model group for interrogating host-virus dynamics due to their limited but well-characterised functional and taxonomic diversity. In this study, we leveraged knowledge of the contrasting ecophysiology between different nitrifier host groups, using urea-stimulated nitrification together with established differential inhibitors to selectively enrich for viruses infecting AOM and NOB in soil microcosms.

One of the major challenges in metagenome-based soil virus ecology is identifying which taxa are infected by individual viruses [90], with the proportion of hosts predicted for viruses in terrestrial system surveys typically lower than that for other environments [68]. Alignment matches between CRISPR spacers and virus genomes provide high confidence linkage predictions but can be challenging in soil samples containing high levels of richness [91]. We previously used ^13^C methane-enriched soils combined with DNA stable-isotope probing to reduce background diversity in metagenomes to focus on methylotroph host-virus interactions. These analyses demonstrated that viruses possessing protospacer sequences matching CRISPR spacers of *Methylocystaceae* could also be linked using a shared homolog approach only [42], which was adopted in this study. The virus prediction tool iPHoP [68] predicted only a total of 13 vOTUs to infect nitrifier hosts and did not confidently identify a nitrifier (or any) host for the majority of nitrifier-infecting vOTUs predicted here using a curated homolog-based analysis. However, further support for the this approach was provided by three different approaches: 1) the majority of AOA and AOB vOTUs were subsequently placed in gene-sharing networks with reference viruses using a custom database of nitrifier proviruses and IMG/VR viruses; 2) one-third of vOTUs also possessed homologs of nitrifier virus-specific hallmark genes, and; 3) the relative abundance of AOA, AOB and *Nitrospira* groups correlated significantly (*p* = ≤0.009) with the relative abundance of their predicted infecting viruses in a series of microcosms under differential inhibition conditions. Compared to AOA and AOB, vOTUs predicted to infect *Nitrospira* strains shared less genetic content with reference viruses. This may indicate that our approach was less successful in identifying *Nitrospira*-infecting vOTUs, but the reduced number of *Nitrospira*-infecting virus genomes (201) in our database compared to those for AOA (1,808) and AOB (1,043) is likely to influence the success of finding matches with reference sequences.

As soil is a largely oligotrophic environment that experiences frequently changing conditions, it has been hypothesised that a high proportion of viruses possess lysogenic capability to enhance survival [22]. However, recent studies indicate that the majority of free soil viruses are not lysogenic [91]. Our previous work found that the majority (86%) of predicted AOA vOTUs were lysogenic but this analysis used untargeted total metagenomes and genes from integrated AOA proviruses in the workflow for predicting AOA-infecting vOTUs, potentially biasing the analysis towards free temperate or host-integrated viruses. In this study, analysis of free virus-targeted and high-quality/complete vOTU genomes only predicted that less than half (33-46%) were capable of lysogeny for all nitrifier groups.

Nitrifier-virus interactions appear to be dynamic in soil and multiple families of viruses were predicted to infect each functional group analysed. It should be noted, however, that there are typically no obvious indications of virus infection influencing nitrification activity or population dynamics in incubation studies like those performed here. Nitrifying soil microcosms usually demonstrate approximately linear increases in net NO_3_^-^ production when NH_4_^+^ is not limiting (e.g. [40,92,93]). Despite being functionally restricted and taxonomically coherent, a range of ecophysiologies exist within ammonia and nitrite oxidiser communities for an individual soil and under a specific set of incubation conditions (e.g. controlled water content, temperature, NH_3_ source, and concentration) only some populations are selected and contribute to nitrification activity. Decreases in the abundance of growing soil nitrifier populations, or variation in N flux rates that could be attributed to viral predation via ‘kill-the-winner’ dynamics, are not typically observed i.e. populations that are selected at the onset of a specific incubation condition invariably continue to grow and dominate activity (e.g. [40,94,95]). This could be a consequence of not observing virus-host interactions with appropriate resolution i.e. while soil nitrifier studies typically characterise individual ‘populations’ from unique 16S rRNA or *amoA* gene sequences, individual ASVs likely represent multiple strains and ‘kill-the-winner’ dynamics may occur within sub-populations below an ASV threshold. An alternative explanation is that nitrifier-virus interactions follow the recently proposed ‘cull-the-winner’ model [91] whereby only a fraction of ‘successful’ growing cells are killed by viral lysis without decimating the entire population and enabling continued growth and contribution to activity. This could even represent a possible explanation why linear (rather than exponential) increases of nitrate and cell numbers are typically observed in soil under incubation conditions with non-limiting substrate.

Soil and marine AOA viruses may relieve different metabolic bottlenecks during infection. Ammonia monooxygenase (AMO) catalyses the first step of the ammonia oxidation pathway in AOM by oxidising NH_3_ to hydroxylamine. In AOA, AMO is encoded by six genes (*amoA, -B, -C, -X, -Y, -Z)* [96] with AOA (and AOB) genomes often containing isolated *amoC* genes in addition to those in gene clusters encoding some or all protein sub-units [35]. Similarly, marine AOA viruses also contain isolated *amoC* genes only [74,97,98] and functionally and evolutionarily related *pmoC* genes are found in viruses infecting methanotrophic populations [42,99]. However, in this and our previous study examining AOA virus diversity in soil [23], *amoC* genes were conspicuously absent, and genes encoding MCO1 were found on 3 of 101 vOTUs from two geographically distant agricultural soils. The enrichment of virome reads mapped to MCO1 compared to other core metabolic genes provided further evidence for the potential importance of virus-encoded MCO1. AOA genomes contain multiple copper (Cu)-binding periplasmic proteins with AOA using Cu in redox reactions during electron transport [100]. While the gene MCO1 was demonstrated to be moderately upregulated in the soil isolate *Nitrososphaera viennensis* under conditions of copper limitation [88], its role in AOA physiology and its potential metabolic role during infection is unclear. However, its identification in multiple virus genomes, and the presence of oxidoreductase genes in other AOA vOTUs, indicate that AOA viruses infection may alleviate bottlenecks in electron transport during infection.

In summary, the characterisation of high-quality genomes of viruses infecting nitrifiers from ‘bulk’ viromes is limited due to the high diversity and complexity of the soil microbiome, together with the low relative abundance of their host populations. These results demonstrate that the use of targeted incubation conditions facilitates the enrichment and recovery of viruses associated with a specific function within the complex soil environment. Future work using similar incubation-based approaches for soils representing a wide range of land-use types and physicochemical properties could facilitate the establishment of a taxonomic framework for nitrifier viruses that is linked to host taxonomy and ecophysiology. As there is considerable interest in inhibiting nitrification activity in agricultural soils, the cultivation and application of nitrifier lytic viruses may be a useful approach for reducing AOM activity after fertilisation events.

## Data availability

All assembled contigs ≥10 kb are available under NCBI BioProject accession number PRJNA1030982 with numerical identifiers as presented in the manuscript. 16S rRNA gene amplicon sequence data are deposited in NCBI’s Sequence Read Archive with accession number PRJNA1010125. Supplementary tables are available as a Google document at https://tinyurl.com/LeeSupplementaryTables.

## Acknowledgments

This work was funded by the Agence Nationale de la Recherche grant ‘CONSERVE’ (ANR-22-CE02-0006-01) awarded to GWN and CH. We acknowledge use of the Newton high-performance computing cluster at PMCS2I, Ećole Centrale de Lyon. The authors would like to thank the Centre de Formation et de Promotion Horticole for access to the l’Abbé Rozier farm and Mr Michael McGibbon (University of Aberdeen) for kindly determining total soil C and N content.

---

**Supplementary Fig. 1.**
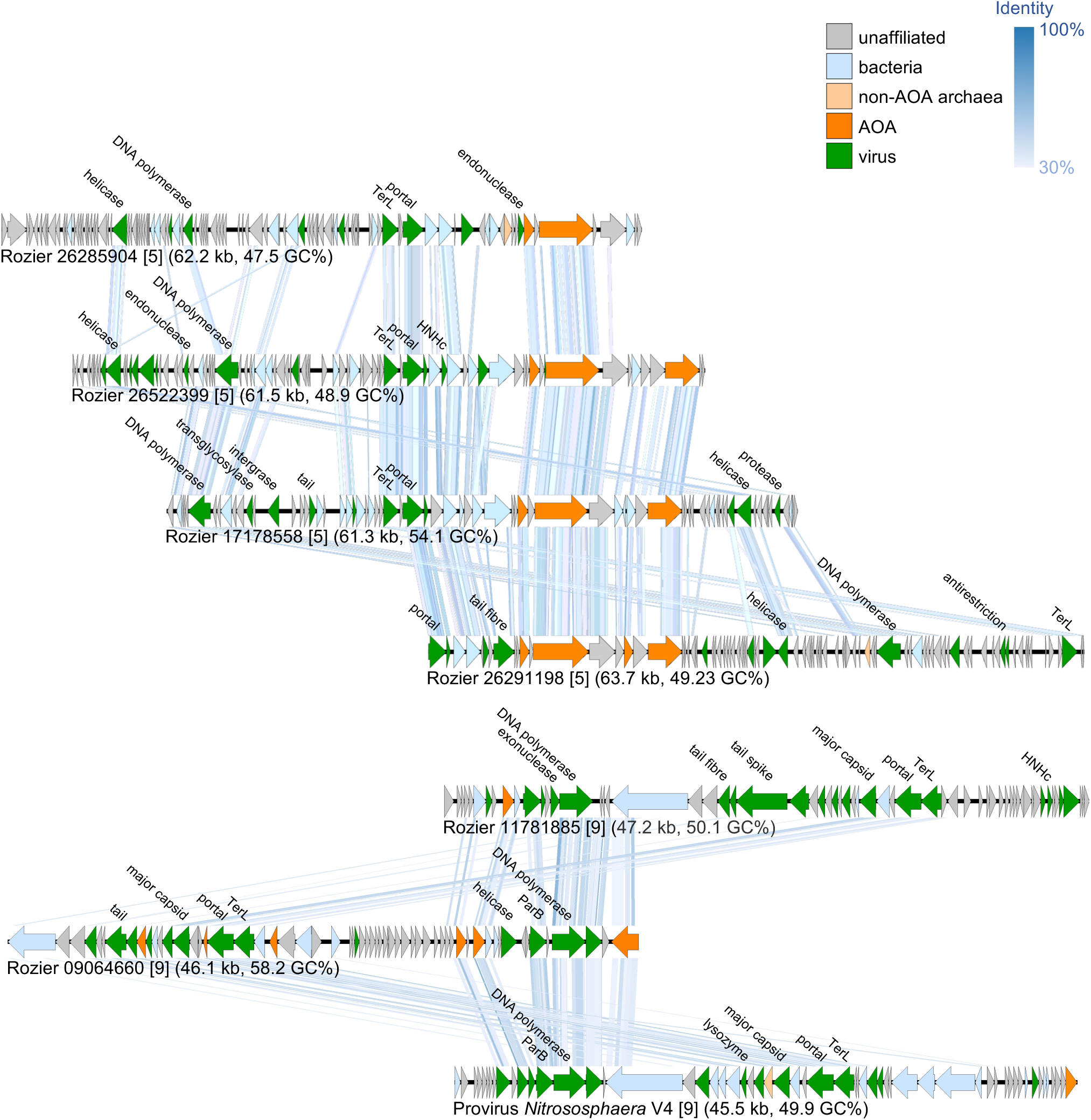
Relative abundance of AOA 16S rRNA gene ASVs amplified from total genomic DNA extracted from soil microcosms incubated with urea and differential inhibitors. Designations were determined using the database of Wang et al. [47] which enables linkage to the *amoA*-gene framework of Alves et al. [5]. Comparisons were made between the same taxonomic group in different samples with different letters indicating significant differences (*p* < 0.05, two-sample Student’s t-test or Welsch’s t-test when variances were not homogenous).

**Supplementary Fig. 2.**
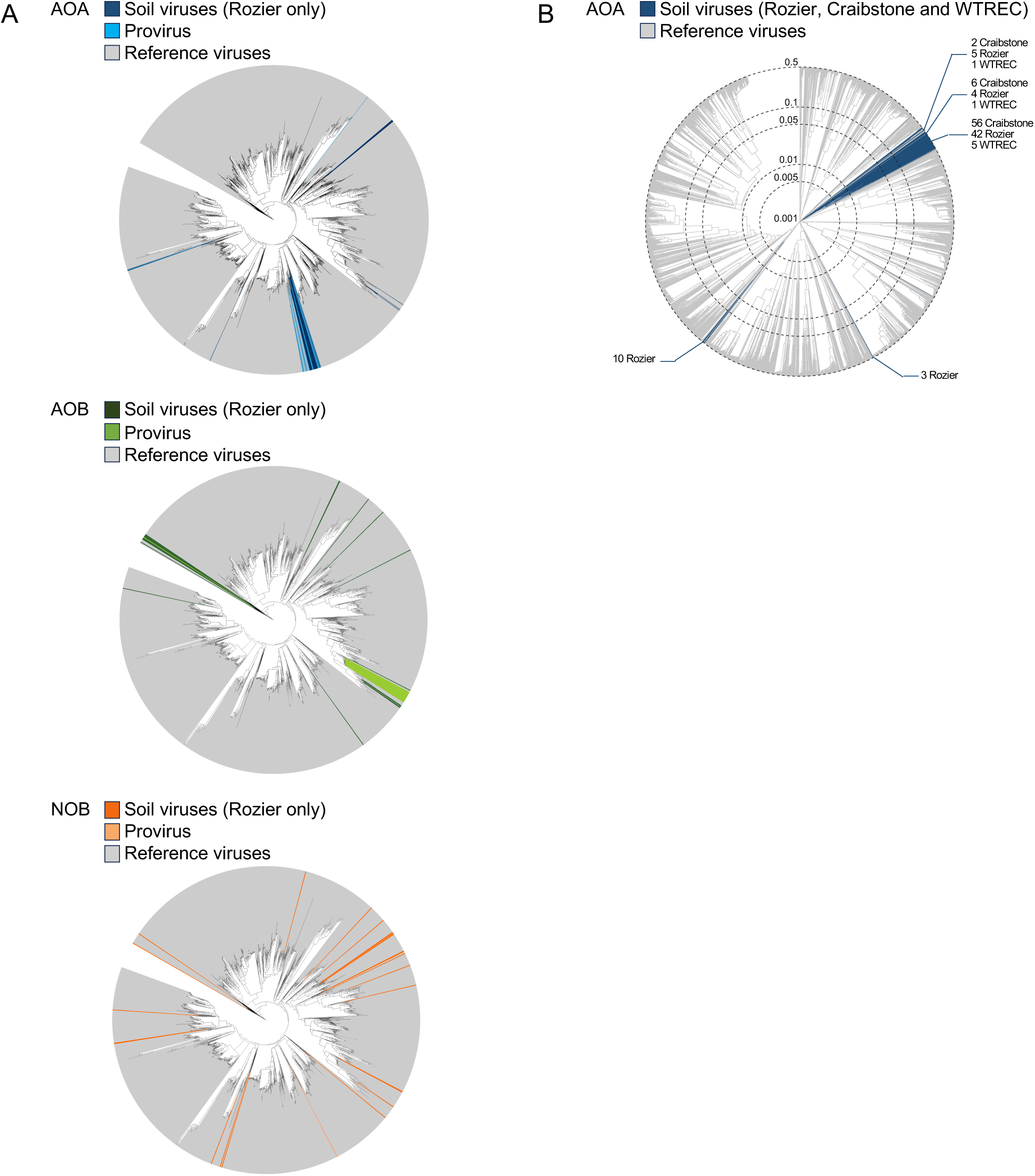
Genome maps of complete or high-quality AOA virus genomes belonging to putative families containing more than one representative. Virus contig names and values in square brackets describing different putative values follow those shown in Fig. 2B. Genes with a predicted virus-specific function are annotated. For full annotations refer to Table S5.

**Supplementary Fig. 3.**
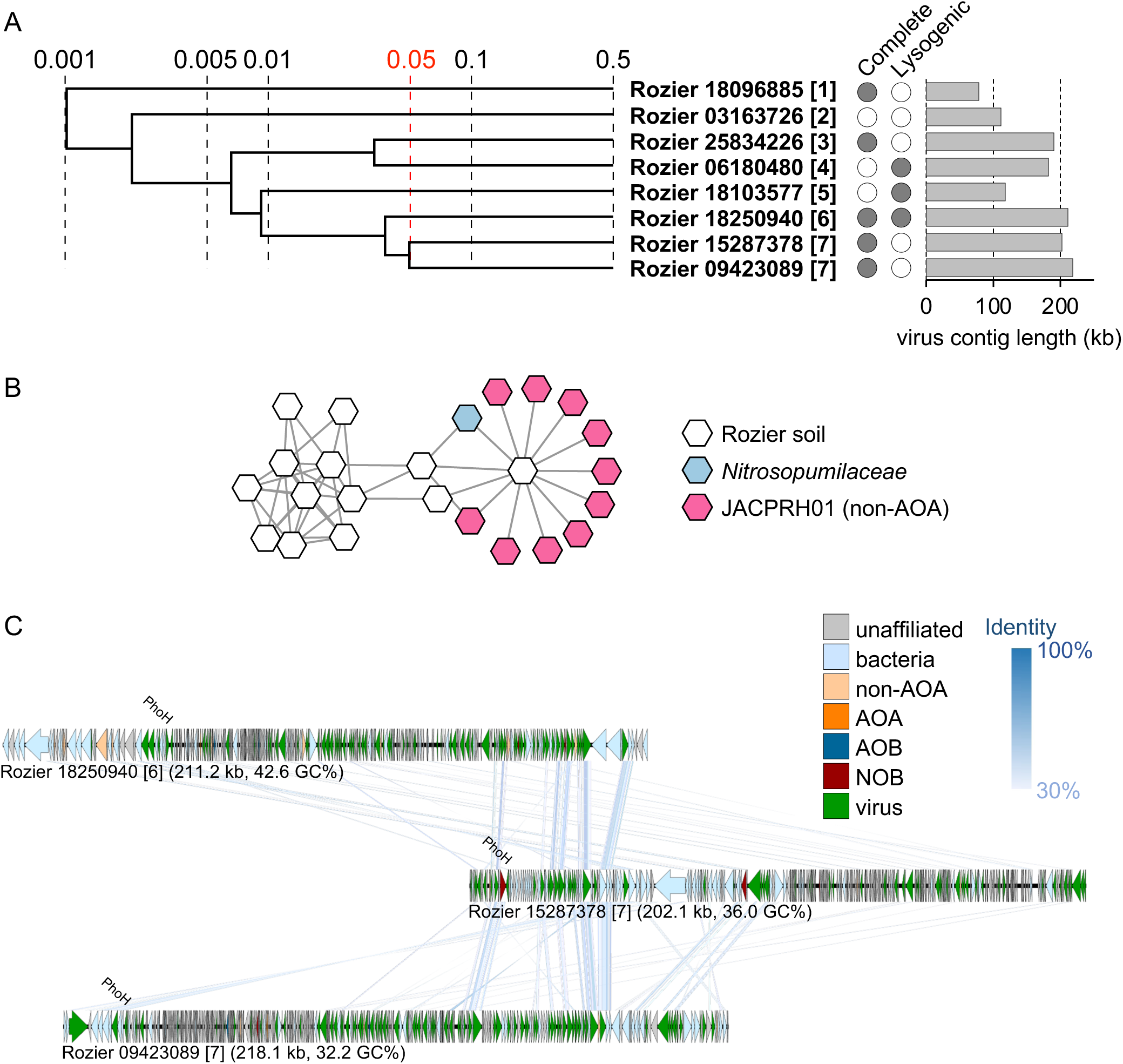
Comparison of AOA, AOB and NOB virus diversity with reference databases. **A** Phylogenetic analysis of derived TerL protein sequences from Rozier soil viruses and nitrifier proviruses together with 1,573 sequences from RefSeq virus genomes. **B** Proteomic tree showing genome-wide sequence similarities between soil AOA virus contigs (≥10 kb) identified in Rozier samples together with those identified in Craibstone, Scotland [23] and the West Tennessee Research and Education Center, USA (WTREC) soils [85]. Reference sequences (n = 5,583) were obtained from the Host-Virus DB [73].

**Supplementary Fig. 4.**
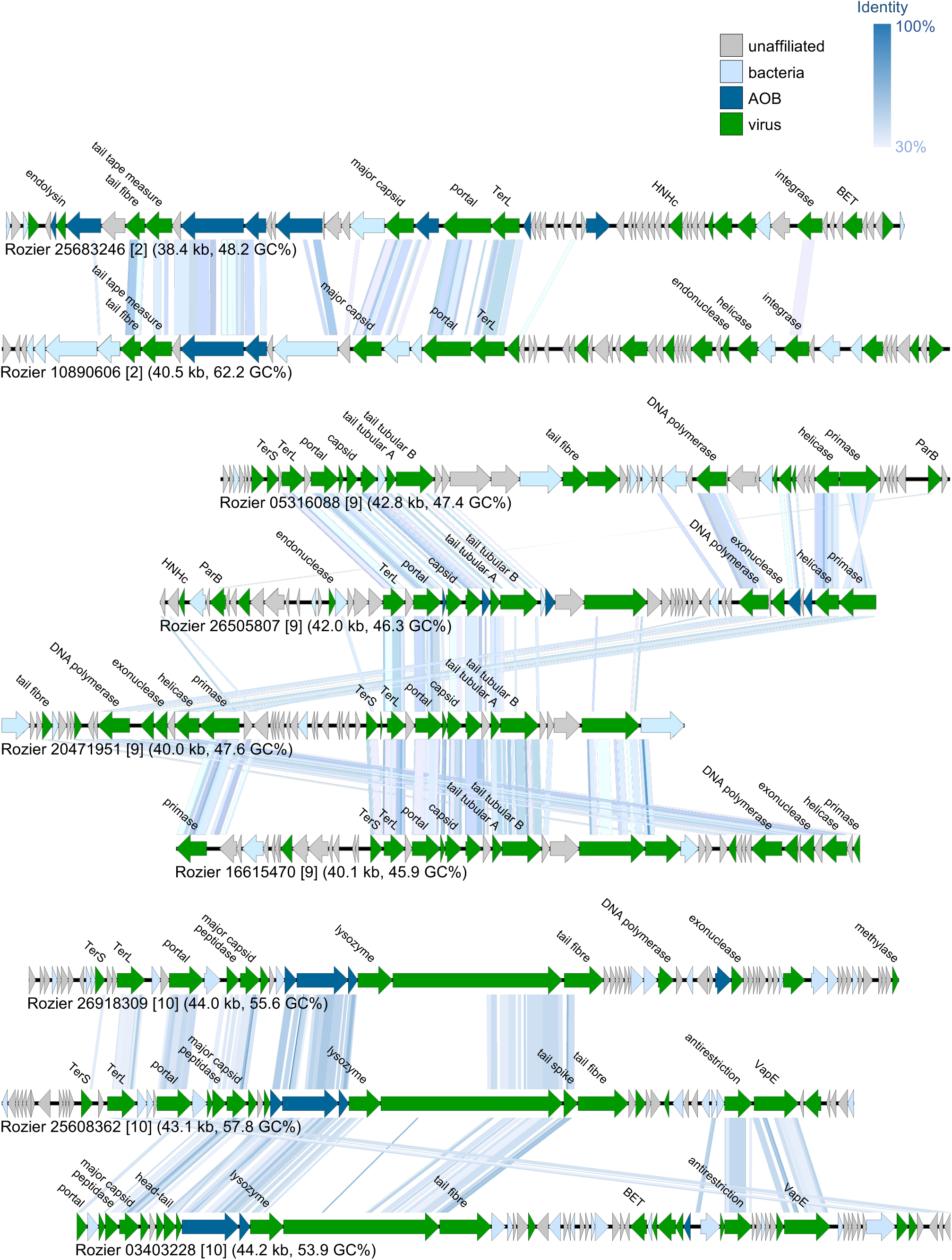
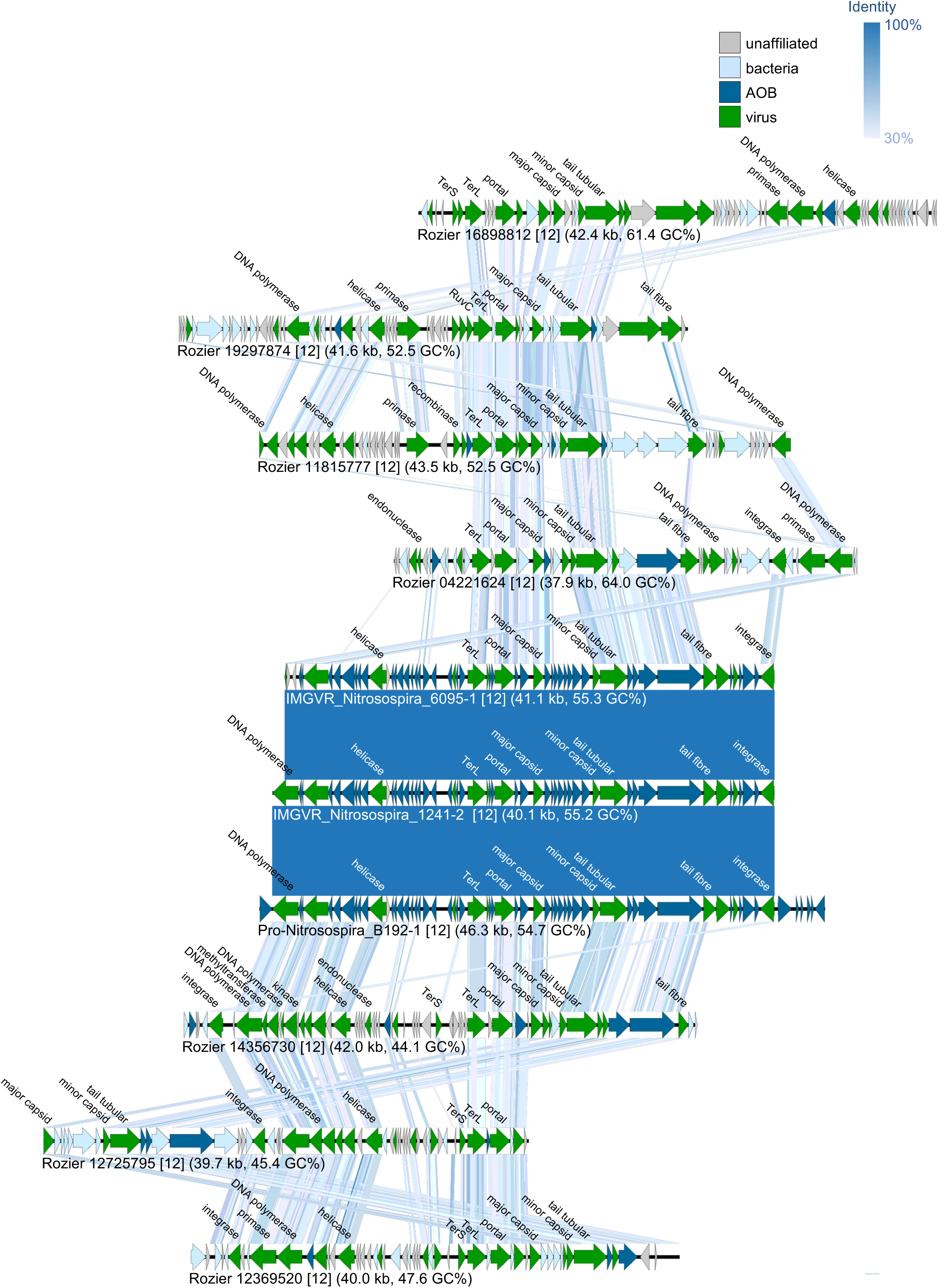
Complete or high-quality virus genomes with homologs shared with non-AOA archaeal genomes. **A** Proteomic tree showing genome-wide sequence similarities between eight high quantity or complete viral contigs only from Rozier soil (name in bold with 8-figure number representing the NCBI contig name). Genome completeness and prediction of lysogeny is denoted by filled circles. Values at dotted lines represent a distance metric based on normalized tBLASTx scores with 0.05 (in red) an estimated threshold for grouping viruses within the same family. Each number in a square bracket denotes an individual putative virus family. **B** Gene-sharing network analysis of contigs ≥10 kb linked with *Nitrososphaera* reference virus sequences. **C** Genome maps of three virus genomes >200 kb placed within two different putative families ([6] and [7] in panel A). For complete annotations refer to Supplementary Table 5.

**Supplementary Fig. 5.**
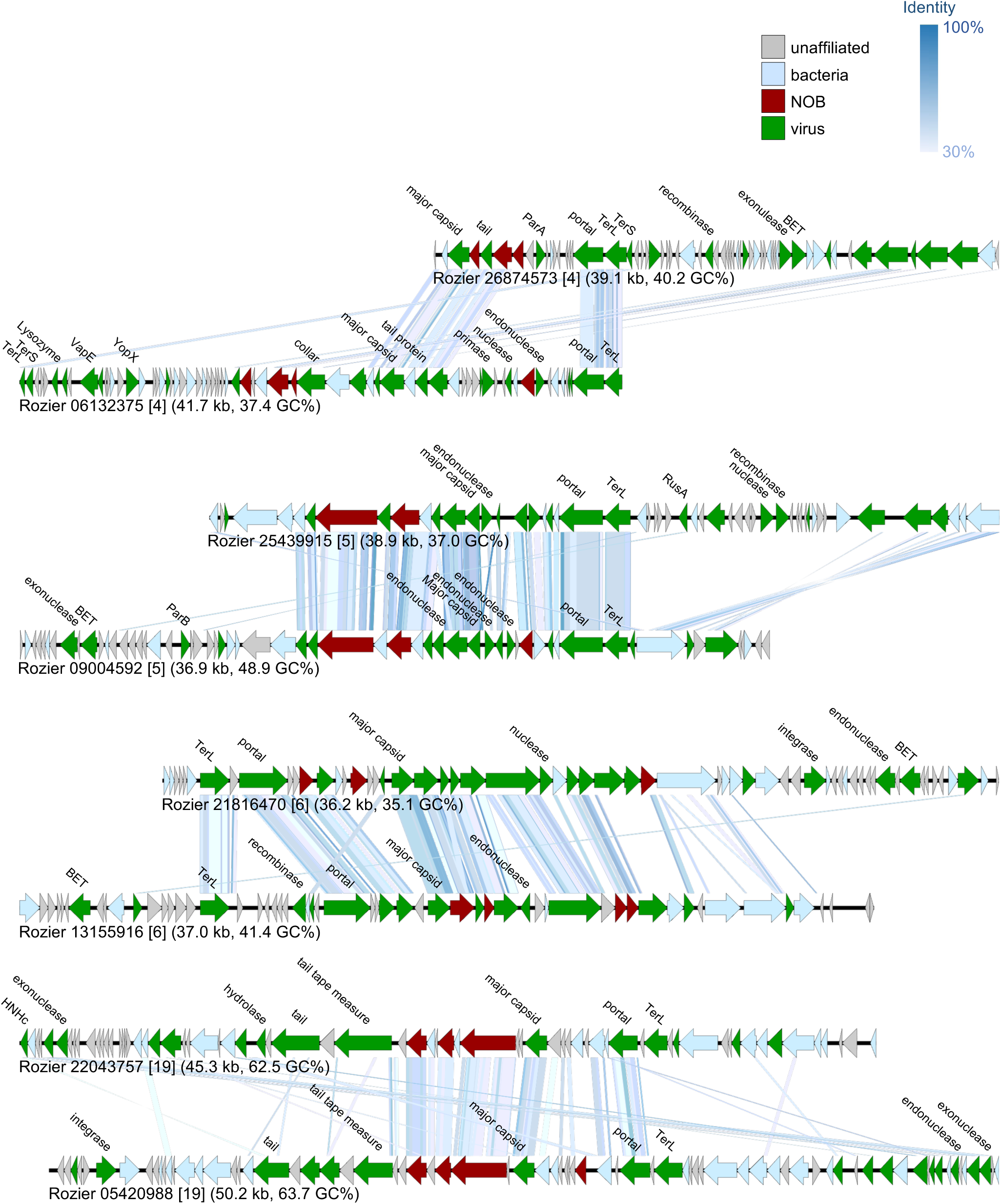
Genome maps of complete or high-quality AOB virus genomes belonging to putative families containing more than one representative. Virus contig names and values in square brackets describing different putative values follow those shown in Fig. 3B. Genes with a predicted virus-specific function are annotated. For full annotations refer to Table S5.

**Supplementary Fig. 6.**
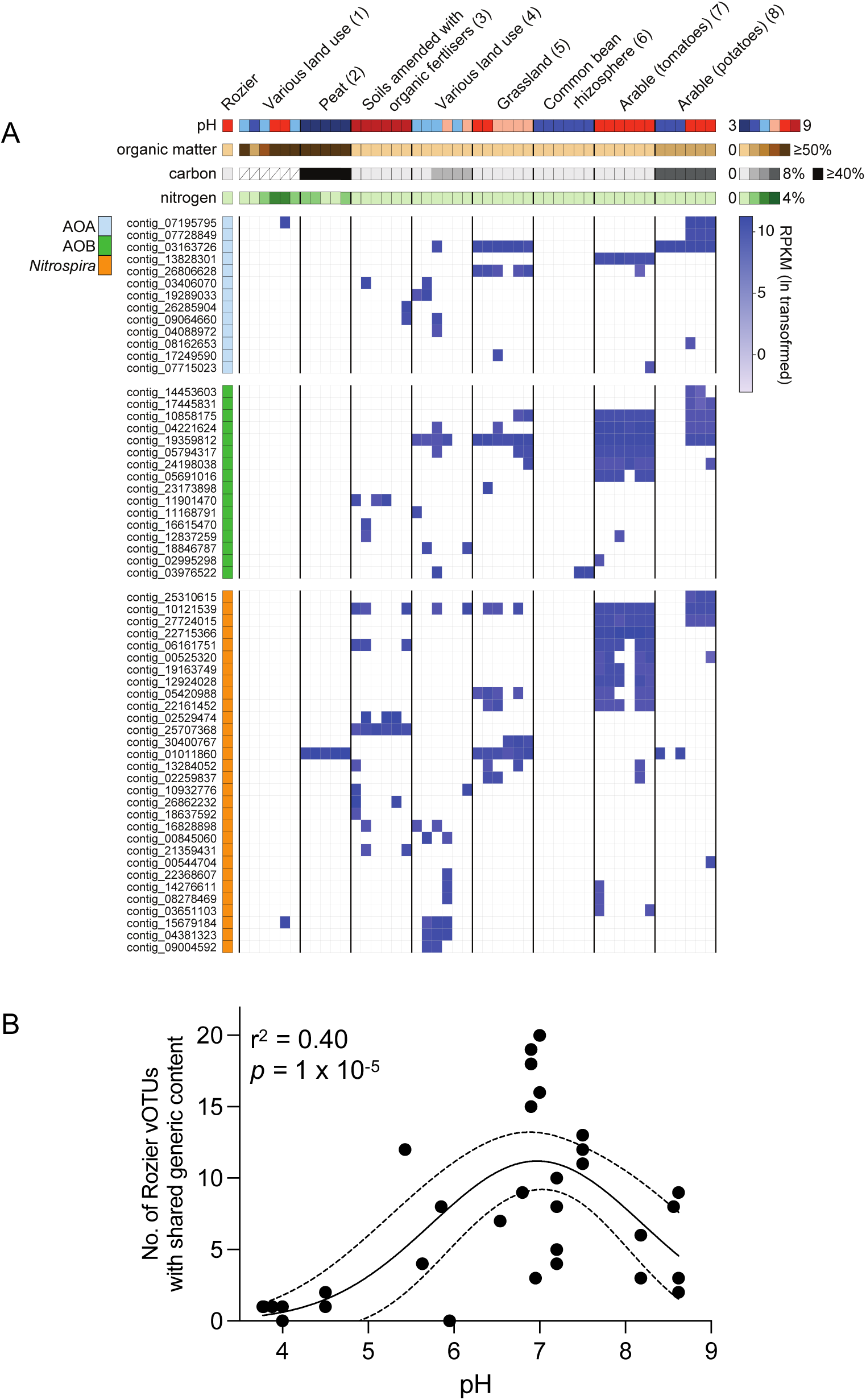
Genome maps of complete or high-quality NOB virus genomes belonging to putative families containing more than one representative. Virus contig names and values in square brackets describing different putative values follow those shown in Fig. 4B. Genes with a predicted virus-specific function are annotated. For full annotations refer to Table S5.

**Supplementary Fig. 7.**
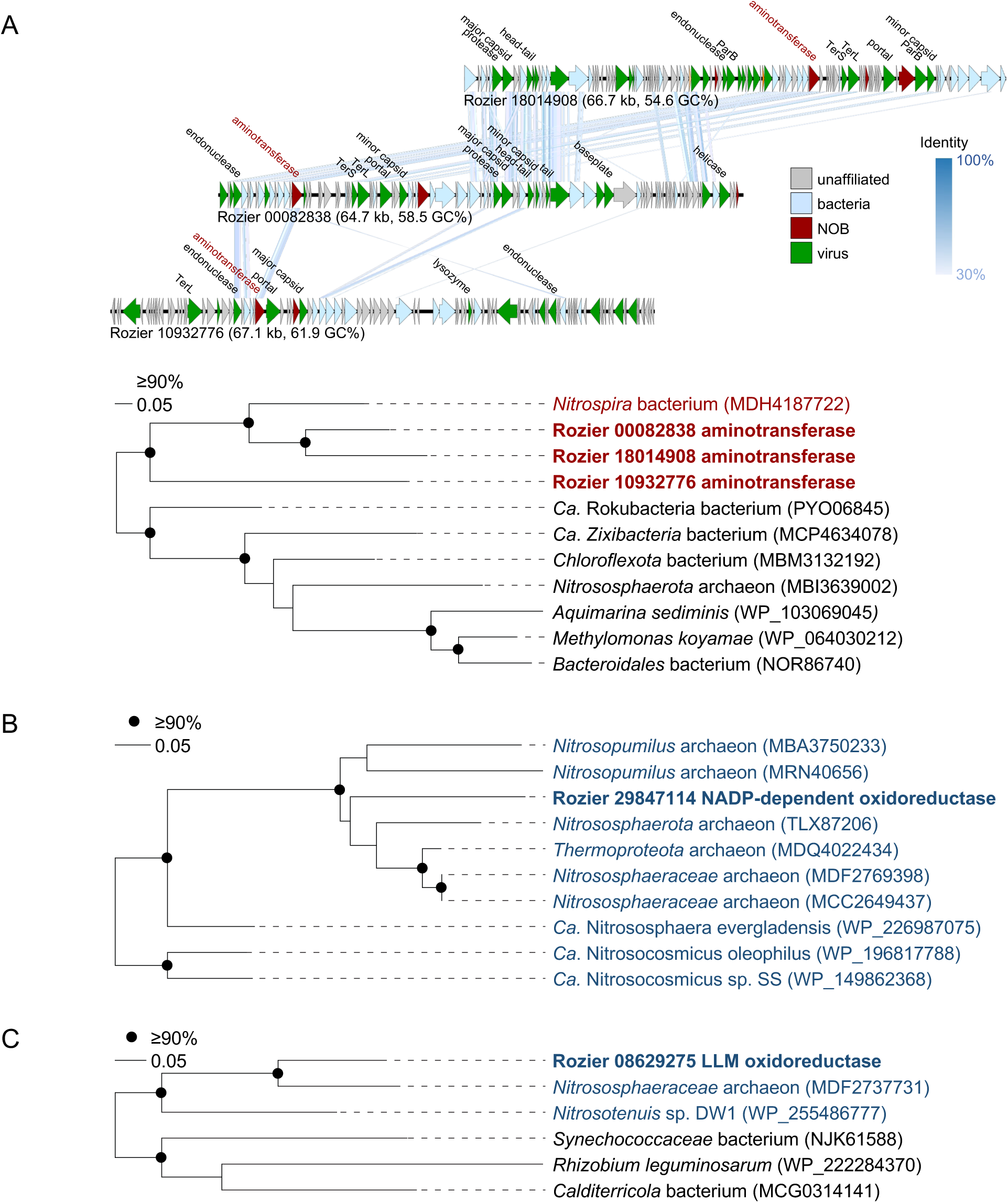
Rozier vOTUs sharing genetic content with selected soil viromes. **A** Rozier vOTUs mapped with virome sequence reads from 41 soil samples in seven studies using a minimum threshold of 10% coverage at 95% identity (relative abundance given as ln-transformed RPKM). Soil physicochemical properties are given within coloured ranges for each soil sample where data is available (pH in 1 unit increments, organic matter content in 10% increments, total C in 2.5% increments, total N in 1% increments). See Table S4 for specific details of each soil sample and derived viromes. **B** Quadratic regression analysis describing number of Rozier vOTUs sharing genetic content with individual soil viromes as a function of soil pH.

**Fig. S8.**
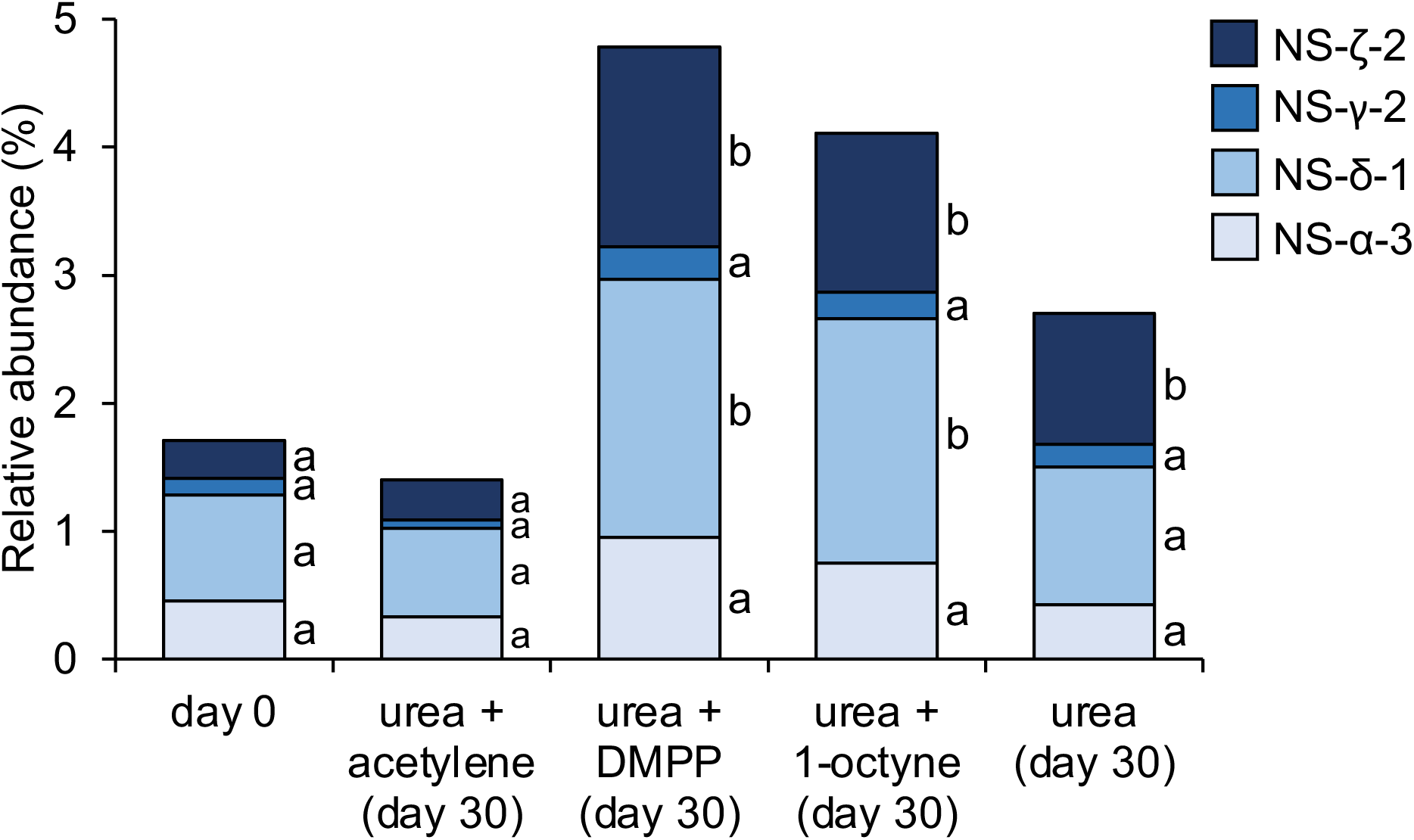
Selected putative auxiliary metabolic genes associated with nitrifying populations. **A** Genome map of three related viruses infecting *Nitrospira* hosts and maximum-likelihood phylogenetic analysis of predicted aminotransferase protein sequences (334 aligned amino acid positions, LG substitution with gamma-distributed sites) with reference sequences from *Nitrospira* and other bacterial strains. **B** Maximum-likelihood phylogenetic analysis of a predicted NADP-dependent oxidoreductase protein sequence found on an AOA virus-associated contig together with those from *Nitrososphaerales* reference genomes (221 aligned positions, Q.pfam substitution and gamma-distributed sites). **C** Maximum-likelihood phylogenetic analysis of a predicted LLM-dependent oxidoreductase protein sequence found on an AOA-infecting virus genome together with those from *Nitrososphaerales* reference genomes (292 aligned positions, Q.plant substitution and gamma-distributed sites).

## References

1. Prosser JI. Autotrophic Nitrification in Bacteria. Advances in Microbial Physiology. Elsevier, 1990, 125–81.

2. Nardi P, Laanbroek HJ, Nicol GW et al. Biological nitrification inhibition in the rhizosphere: determining interactions and impact on microbially mediated processes and potential applications. FEMS Microbiol Rev 2020;44:874–908.

3. Lassaletta L, Billen G, Grizzetti B et al. 50 year trends in nitrogen use efficiency of world cropping systems: the relationship between yield and nitrogen input to cropland. Environ Res Lett 2014;9:105011.

4. Prosser JI, Hink L, Gubry-Rangin C et al. Nitrous oxide production by ammonia oxidizers: Physiological diversity, niche differentiation and potential mitigation strategies. Glob Chang Biol 2020;26:103–18.

5. Alves RJE, Eloy Alves RJ, Minh BQ et al. Unifying the global phylogeny and environmental distribution of ammonia-oxidising archaea based on amoA genes. Nature Communications 2018;9:1517.

6. Li Y, Chapman SJ, Nicol GW et al. Nitrification and nitrifiers in acidic soils. Soil Biol Biochem 2018;116:290–301.

7. Daims H, Lebedeva EV, Pjevac P et al. Complete nitrification by *Nitrospira* bacteria. Nature 2015;528:504–9.

8. van Kessel MAHJ, Speth DR, Albertsen M et al. Complete nitrification by a single microorganism. Nature 2015;528:555–9.

9. Daims H, Lücker S, Wagner M. A New Perspective on Microbes Formerly Known as Nitrite-Oxidizing Bacteria. Trends Microbiol 2016;24:699–712.

10. Prosser JI, Nicol GW. Relative contributions of archaea and bacteria to aerobic ammonia oxidation in the environment. Environ Microbiol 2008;10:2931–41.

11. Hink L, Nicol GW, Prosser JI. Archaea produce lower yields of N2 O than bacteria during aerobic ammonia oxidation in soil. Environ Microbiol 2017;19:4829–37.

12. Hink L, Gubry-Rangin C, Nicol GW et al. The consequences of niche and physiological differentiation of archaeal and bacterial ammonia oxidisers for nitrous oxide emissions. ISME J 2018;12:1084–93.

13. Hazard C, Prosser JI, Nicol GW. Use and abuse of potential rates in soil microbiology. Soil Biol Biochem 2021;157:108242.

14. Taylor AE, Giguere AT, Zoebelein CM et al. Modeling of soil nitrification responses to temperature reveals thermodynamic differences between ammonia-oxidizing activity of archaea and bacteria. ISME J 2017;11:896–908.

15. Killham K. Nitrification in coniferous forest soils. Plant Soil 1990;128:31–44.

16. Taylor AE, Vajrala N, Giguere AT et al. Use of aliphatic n-alkynes to discriminate soil nitrification activities of ammonia-oxidizing thaumarchaea and bacteria. Appl Environ Microbiol 2013;79:6544–51.

17. Duan P, Wu Z, Zhang Q et al. Thermodynamic responses of ammonia-oxidizing archaea and bacteria explain N2O production from greenhouse vegetable soils. Soil Biol Biochem 2018;120:37–47.

18. Papadopoulou ES, Bachtsevani E, Lampronikou E et al. Comparison of novel and established nitrification inhibitors relevant to agriculture on soil ammonia- and nitrite-oxidizing isolates. Front Microbiol 2020;11:581283.

19. Shen T, Stieglmeier M, Dai J et al. Responses of the terrestrial ammonia-oxidizing archaeon Ca. *Nitrososphaera viennensis* and the ammonia-oxidizing bacterium *Nitrosospira multiformis* to nitrification inhibitors. FEMS Microbiol Lett 2013;344:121–9.

20. Lehtovirta-Morley LE, Verhamme DT, Nicol GW et al. Effect of nitrification inhibitors on the growth and activity of *Nitrosotalea devanaterra* in culture and soil. Soil Biol Biochem 2013;62:129–33.

21. Braga LPP, Spor A, Kot W et al. Impact of phages on soil bacterial communities and nitrogen availability under different assembly scenarios. Microbiome 2020;8:52.

22. Roux S, Emerson JB. Diversity in the soil virosphere: to infinity and beyond? Trends Microbiol 2022;30:1025–35.

23. Lee S, Sieradzki ET, Nicol GW et al. Propagation of viral genomes by replicating ammonia-oxidising archaea during soil nitrification. ISME J 2023;17:309–14.

24. Klein T, Poghosyan L, Barclay JE et al. Cultivation of ammonia-oxidising archaea on solid medium. FEMS Microbiol Lett 2022;369:fnac029.

25. Bock E, Sundermeyer-Klinger H, Stackebrandt E. New facultative lithoautotrophic nitrite-oxidizing bacteria. Arch Microbiol 1983;136:281–4.

26. Koops HP, Böttcher B, Möller UC et al. Classification of eight new species of ammonia-oxidizing bacteria: *Nitrosomonas communis* sp. nov., *Nitrosomonas ureae* sp. nov., *Nitrosomonas aestuarii* sp. nov., *Nitrosomonas marina* sp. nov., *Nitrosomonas nitrosa* sp. nov., *Nitrosomonas eutropha* sp. nov., Nitrosomonas oligotropha sp. nov. and Nitrosomonas halophila sp. nov. J Gen Microbiol 1991;137:1689–99.

27. Krupovic M, Quemin ERJ, Bamford DH et al. Unification of the globally distributed spindle-shaped viruses of the Archaea. J Virol 2014;88:2354–8.

28. Kim J-G, Kim S-J, Cvirkaite-Krupovic V et al. Spindle-shaped viruses infect marine ammonia-oxidizing thaumarchaea. Proc Natl Acad Sci U S A 2019;116:15645–50.

29. Quirós P, Sala-Comorera L, Gómez-Gómez C et al. Identification of a virulent phage infecting species of *Nitrosomonas*. ISME J 2023;17:645–8.

30. Chain P, Lamerdin J, Larimer F et al. Complete genome sequence of the ammonia-oxidizing bacterium and obligate chemolithoautotroph *Nitrosomonas europaea*. J Bacteriol 2003;185:2759–73.

31. Stein LY, Arp DJ, Berube PM et al. Whole-genome analysis of the ammonia-oxidizing bacterium, *Nitrosomonas eutropha* C91: implications for niche adaptation. Environ Microbiol 2007;9:2993–3007.

32. Choi J, Kotay SM, Goel R. Various physico-chemical stress factors cause prophage induction in *Nitrosospira multiformis* 25196--an ammonia oxidizing bacteria. Water Res 2010;44:4550–8.

33. Krupovic M, Spang A, Gribaldo S et al. A thaumarchaeal provirus testifies for an ancient association of tailed viruses with archaea. Biochem Soc Trans 2011;39:82–8.

34. Abby SS, Melcher M, Kerou M et al. *Candidatus* Nitrosocaldus cavascurensis, an ammonia oxidizing, extremely thermophilic archaeon with a highly mobile genome. Front Microbiol 2018;9:28.

35. Spang A, Poehlein A, Offre P et al. The genome of the ammonia-oxidizing *Candidatus* Nitrososphaera gargensis: insights into metabolic versatility and environmental adaptations. Environ Microbiol 2012;14:3122–45.

36. Rice MC, Norton JM, Valois F et al. Complete genome of *Nitrosospira briensis* C-128, an ammonia-oxidizing bacterium from agricultural soil. Stand Genomic Sci 2016;11:46.

37. Ushiki N, Fujitani H, Shimada Y et al. Genomic analysis of two phylogenetically distinct *Nitrospira* species reveals their genomic plasticity and functional diversity. Front Microbiol 2018;8:fmicb.2017.02637.

38. Payne LJ, Meaden S, Mestre MR et al. PADLOC: a web server for the identification of antiviral defence systems in microbial genomes. Nucleic Acids Res 2022;50:W541–50.

39. Bachtsevani E, Papazlatani CV, Rousidou C et al. Effects of the nitrification inhibitor 3,4-dimethylpyrazole phosphate (DMPP) on the activity and diversity of the soil microbial community under contrasting soil pH. Biol Fertil Soils 2021;57:1117–35.

40. Offre P, Prosser JI, Nicol GW. Growth of ammonia-oxidizing archaea in soil microcosms is inhibited by acetylene. FEMS Microbiol Ecol 2009;70:99–108.

41. Trubl G, Solonenko N, Chittick L et al. Optimization of viral resuspension methods for carbon-rich soils along a permafrost thaw gradient. PeerJ 2016;4:e1999.

42. Lee S, Sieradzki ET, Nicolas AM et al. Methane-derived carbon flows into host– virus networks at different trophic levels in soil. Proceedings of the National Academy of Sciences 2021;118:e2105124118.

43. Nicol GW, Prosser JI. Strategies to determine diversity, growth, and activity of ammonia-oxidizing archaea in soil. Methods Enzymol 2011;496:3–34.

44. Walters W, Hyde ER, Berg-Lyons D et al. Improved bacterial 16S rRNA gene (V4 and V4-5) and fungal internal transcribed spacer marker gene primers for microbial community surveys. mSystems 2016;1:e00009–15.

45. Callahan BJ, McMurdie PJ, Rosen MJ et al. DADA2: High-resolution sample inference from Illumina amplicon data. Nat Methods 2016;13:581–3.

46. Quast C, Pruesse E, Yilmaz P et al. The SILVA ribosomal RNA gene database project: improved data processing and web-based tools. Nucleic Acids Res 2013;41:D590–6.

47. Wang H, Bagnoud A, Ponce-Toledo RI et al. Linking 16S rRNA gene classification to *amoA* gene taxonomy reveals environmental distribution of ammonia-oxidizing Archaeal clades in peatland soils. mSystems 2021;6:00546–21.

48. Uritskiy GV, DiRuggiero J, Taylor J. MetaWRAP—a flexible pipeline for genome-resolved metagenomic data analysis. Microbiome 2018;6:1–13.

49. Lee S, Sorensen JW, Walker RL et al. Soil pH influences the structure of virus communities at local and global scales. Soil Biol Biochem 2022;166:108569.

50. Li D, Liu C-M, Luo R et al. MEGAHIT: an ultra-fast single-node solution for large and complex metagenomics assembly via succinct de Bruijn graph. Bioinformatics 2015;31:1674–6.

51. Roux S, Enault F, Hurwitz BL et al. VirSorter: mining viral signal from microbial genomic data. PeerJ 2015;3:e985.

52. Guo J, Bolduc B, Zayed AA et al. VirSorter2: a multi-classifier, expert-guided approach to detect diverse DNA and RNA viruses. Microbiome 2021;9:37.

53. Ren J, Song K, Deng C et al. Identifying viruses from metagenomic data using deep learning. Quant Biol 2020;8:64–77.

54. Camargo AP, Roux S, Schulz F et al. Identification of mobile genetic elements with geNomad. Nat Biotechnol 2023;10.1038/s41587-023-01953-y.

55. Nayfach S, Camargo AP, Schulz F et al. CheckV assesses the quality and completeness of metagenome-assembled viral genomes. Nat Biotechnol 2021;39:578–85.

56. Kieft K, Zhou Z, Anantharaman K. VIBRANT: automated recovery, annotation and curation of microbial viruses, and evaluation of viral community function from genomic sequences. Microbiome 2020;8:90.

57. Roux S, Adriaenssens EM, Dutilh BE et al. Minimum Information about an Uncultivated Virus Genome (MIUViG). Nat Biotechnol 2019;37:29–37.

58. Hyatt D, Chen G-L, Locascio PF et al. Prodigal: prokaryotic gene recognition and translation initiation site identification. BMC Bioinformatics 2010;11:119.

59. Buchfink B, Xie C, Huson DH. Fast and sensitive protein alignment using DIAMOND. Nature Methods 2015;12:59–60.

60. Besemer J. GeneMarkS: a self-training method for prediction of gene starts in microbial genomes. Implications for finding sequence motifs in regulatory regions. Nucleic Acids Res 2001;29:2607–18.

61. Suzuki S, Ishida T, Ohue M et al. GHOSTX: A fast sequence homology search tool for functional annotation of metagenomic data. Methods in Molecular Biology. New York, NY: Springer New York, 2017, 15–25.

62. Eddy SR. Accelerated profile HMM searches. PLoS Comput Biol 2011;7:e1002195.

63. Bushnell B. BBMap: A Fast, Accurate, Splice-Aware Aligner. Lawrence Berkeley National Lab. (LBNL), Berkeley, CA (United States), 2014.

64. Imelfort M, Woodcroft B, Parks D. BamM Software package.

65. Kolde R. pheatmap: Pretty Heatmaps. R package version 1.0.12. 2019.

66. Kassambara A. rstatix: Pipe-friendly framework for basic statistical tests. R package version 0.7.2. 2023.

67. Al-Shayeb B, Sachdeva R, Chen L-X et al. Clades of huge phages from across Earth’s ecosystems. Nature 2020;578:425–31.

68. Roux S, Camargo AP, Coutinho FH et al. iPHoP: An integrated machine learning framework to maximize host prediction for metagenome-derived viruses of archaea and bacteria. PLoS Biol 2023;21:e3002083.

69. Parks DH, Chuvochina M, Rinke C et al. GTDB: an ongoing census of bacterial and archaeal diversity through a phylogenetically consistent, rank normalized and complete genome-based taxonomy. Nucleic Acids Res 2022;50:D785–94.

70. Camargo AP, Nayfach S, Chen I-MA et al. IMG/VR v4: an expanded database of uncultivated virus genomes within a framework of extensive functional, taxonomic, and ecological metadata. Nucleic Acids Res 2023;51:D733–43.

71. Sirén K, Millard A, Petersen B et al. Rapid discovery of novel prophages using biological feature engineering and machine learning. bioRxiv 2020; 10.1101/2020.08.09.243022.

72. Bin Jang H, Bolduc B, Zablocki O et al. Taxonomic assignment of uncultivated prokaryotic virus genomes is enabled by gene-sharing networks. Nat Biotechnol 2019;37:632–9.

73. Nishimura Y, Yoshida T, Kuronishi M et al. ViPTree: the viral proteomic tree server. Bioinformatics 2017;33:2379–80.

74. Zhou Y, Zhou L, Yan S et al. Diverse viruses of marine archaea discovered using metagenomics. Environ Microbiol 2023;25:367–82.

75. Sullivan MJ, Petty NK, Beatson SA. Easyfig: a genome comparison visualizer. Bioinformatics 2011;27:1009–10.

76. Brister JR, Ako-Adjei D, Bao Y et al. NCBI viral genomes resource. Nucleic Acids Res 2015;43:D571–7.

77. Katoh K, Standley DM. MAFFT multiple sequence alignment software version 7: improvements in performance and usability. Mol Biol Evol 2013;30:772–80.

78. Capella-Gutiérrez S, Silla-Martínez JM, Gabaldón T. trimAl: a tool for automated alignment trimming in large-scale phylogenetic analyses. Bioinformatics 2009;25:1972–3.

79. Nguyen L-T, Schmidt HA, von Haeseler A et al. IQ-TREE: a fast and effective stochastic algorithm for estimating maximum-likelihood phylogenies. Mol Biol Evol 2015;32:268–74.

80. Edgar RC. MUSCLE: multiple sequence alignment with high accuracy and high throughput. Nucleic Acids Res 2004;32:1792–7.

81. Santos-Medellin C, Zinke LA, Ter Horst AM et al. Viromes outperform total metagenomes in revealing the spatiotemporal patterns of agricultural soil viral communities. ISME J 2021;15:1956–70.

82. Jurgens G, Lindström K, Saano A. Novel group within the kingdom Crenarchaeota from boreal forest soil. Appl Environ Microbiol 1997;63:803–5.

83. Sheridan PO, Meng Y, Williams TA et al. Genomics of soil depth niche partitioning in the Thaumarchaeota family Gagatemarchaeaceae. Nat Commun 2023;14:7305.

84. Cardinale DJ, Duffy S. Single-stranded genomic architecture constrains optimal codon usage. Bacteriophage 2011;1:219–24.

85. Duan N, Radosevich M, Zhuang J et al. Identification of novel viruses and their microbial hosts from soils with long-term nitrogen fertilization and cover cropping management. mSystems 2022;7:e0057122.

86. Yuan Y, Gao M. Jumbo Bacteriophages: An Overview. Front Microbiol 2017;8:fmicb.2017.00403.

87. Ter Horst AM, Santos-Medellín C, Sorensen JW et al. Minnesota peat viromes reveal terrestrial and aquatic niche partitioning for local and global viral populations. Microbiome 2021;9:233.

88. Reyes C, Hodgskiss LH, Kerou M et al. Genome wide transcriptomic analysis of the soil ammonia oxidizing archaeon *Nitrososphaera viennensis* upon exposure to copper limitation. ISME J 2020;14:2659–74.

89. Han L-L, Yu D-T, Bi L et al. Distribution of soil viruses across China and their potential role in phosphorous metabolism. Environ Microbiome 2022;17:6.

90. Coclet C, Roux S. Global overview and major challenges of host prediction methods for uncultivated phages. Curr Opin Virol 2021;49:117–26.

91. Santos-Medellín C, Blazewicz SJ, Pett-Ridge J et al. Viral but not bacterial community successional patterns reflect extreme turnover shortly after rewetting dry soils. Nat Ecol Evol 2023;7:1809–22.

92. Gubry-Rangin C, Nicol GW, Prosser JI. Archaea rather than bacteria control nitrification in two agricultural acidic soils. FEMS Microbiol Ecol 2010;74:566–74.

93. Zhang L-M, Hu H-W, Shen J-P et al. Ammonia-oxidizing archaea have more important role than ammonia-oxidizing bacteria in ammonia oxidation of strongly acidic soils. ISME J 2012;6:1032–45.

94. Webster G, Embley TM, Freitag TE et al. Links between ammonia oxidizer species composition, functional diversity and nitrification kinetics in grassland soils. Environ Microbiol 2005;7:676–84.

95. Tourna M, Freitag TE, Nicol GW et al. Growth, activity and temperature responses of ammonia-oxidizing archaea and bacteria in soil microcosms. Environ Microbiol 2008;10:1357–64.

96. Hodgskiss LH, Melcher M, Kerou M et al. Unexpected complexity of the ammonia monooxygenase in archaea. ISME J 2023;17:588–99.

97. Roux S, Coordinators TO, Brum JR et al. Ecogenomics and potential biogeochemical impacts of globally abundant ocean viruses. Nature 2016;537:689–93.

98. Ahlgren NA, Fuchsman CA, Rocap G et al. Discovery of several novel, widespread, and ecologically distinct marine Thaumarchaeota viruses that encode amoC nitrification genes. ISME J 2019;13:618–31.

99. Chen L-X, Méheust R, Crits-Christoph A et al. Large freshwater phages with the potential to augment aerobic methane oxidation. Nat Microbiol 2020;5:1504–15.

100. Walker CB, de la Torre JR, Klotz MG et al. *Nitrosopumilus maritimus* genome reveals unique mechanisms for nitrification and autotrophy in globally distributed marine crenarchaea. Proc Natl Acad Sci U S A 2010;107:8818–23.

